# A single-cell transcriptome atlas of adult male and female human hookworm *Ancylostoma ceylanicum*

**DOI:** 10.1101/2025.01.03.631133

**Authors:** Suman Bharti, Bruce A Rosa, John Martin, Makedonka Mitreva

**Affiliations:** Department of Medicine, Washington University School of Medicine, St. Louis, Missouri, USA; Department of Genetics, Washington University School of Medicine, St. Louis, Missouri, USA; McDonnell Genome Institute, Washington University in St. Louis, Missouri, USA

**Keywords:** Single-cell transcriptomics, hookworm, parasitology, cellular heterogeneity, tissue annotation, cell atlas

## Abstract

Hookworms infect over 500 million people globally and represent a major neglected tropical disease. The prevalence, high reinfection rate, limited drugs and emerging resistance underscore the need for new therapeutic or vaccine targets. To mitigate halted progress of discovering targets, we provide new molecular insights into sex-specific expression and cellular heterogeneity by generating the first single-cell atlas of male and female transcriptome for the human hookworm *Ancylostoma ceylanicum*. We systemically annotated 10 transcriptionally distinct cell types in female and 13 in male hookworm by comparing marker genes with tissue-specific knowledge. Non-reproductive tissues included pharynx, intestine, cephalic and pharyngeal glands, muscle and hypodermis, complemented with female reproductive tissues (ovary, uterus, eggs, seminal receptacle), and male reproductive tissues (testis, seminal vesicle, cement gland, sperm). Anatomical confocal microscopy and fluorescence in situ hybridization assisted in visualizing these tissues and confirming marker genes. Functional comparisons revealed previously undescribed tissue pathways and molecular markers, providing previously unrecognized insights into hookworm biology, and new targets. Overall, this study presents a detailed cell-type atlas for adult stages of this important yet neglected metazoan parasite.

**Graphical abstract:** 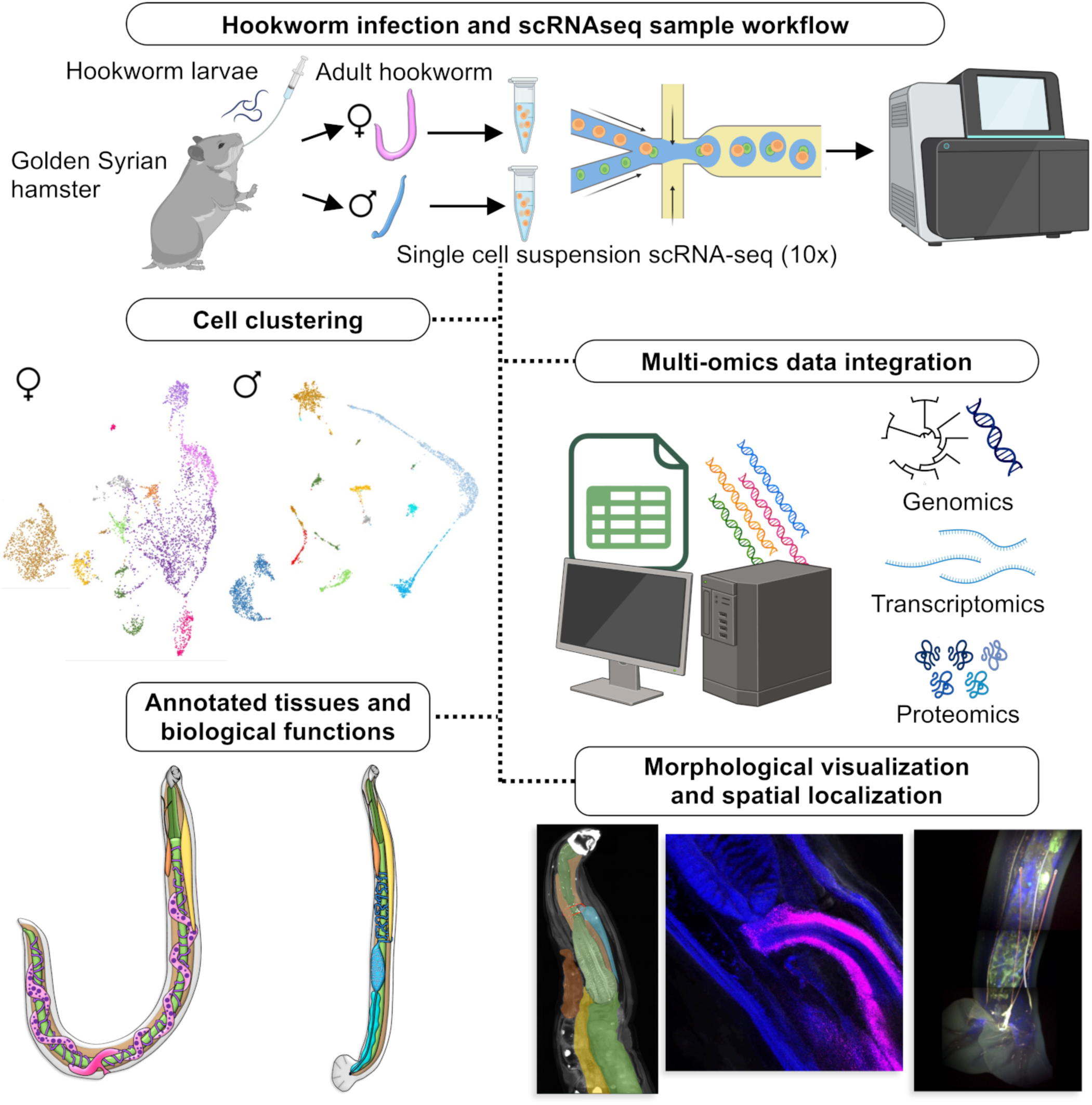

## Introduction

Hookworms are soil transmitted parasitic nematodes that affect an estimated 500 million individuals in the tropical region of the world and result in the loss of approximately 4.1 million disability-adjusted life years (DALYs) annually^1^, representing the most important of the neglected tropical diseases (NTDs). Hookworms can reside in the small intestine for years, and their blood-feeding behavior (hematophagous) can lead to iron-deficiency anemia, malnutrition, stunted growth and development in children, as well as severe morbidity and mortality during pregnancy in women with moderate and heavy infections^1^. It is estimated that each female worm ingests a minimum of 0.1 mL of blood per day^2^, and they survive in the host on average 1-3 years for *Ancylostoma duodenale* and 3-10 years for *Necator americanus*^3^, and their life span can extend up to 18 years)^4^. Hookworm infestation inflicts significant harm on the most vulnerable of the world’s population, including children and women of childbearing age^5^. The world relies on two anthelminthic drugs for deworming, however high prevalence persists in some countries, in part due to high levels of reinfection after treatment^6^ and reports on resistance in some hookworm species^7^ underscores the urgent need for development of preventive and curative interventions or novel strategies to treat and eliminate this NTD.

Given the global importance of these parasites, resources have been developed to enhance the molecular understanding of the three hookworm species infect humans: *Necator americanus*, *A. duodenale* and *A. ceylanicum.* As demonstrated with *N. americanus* following the publication of its genomes^8^, analyzing the hookworm genome provides insights into its molecular complexity and enables the identification and characterization of genes involved in host invasion, blood-feeding, development, and host-parasite interactions, which are crucial for the parasite’s long-term survival within the host. This study was followed by the genome of *A. ceylanicum*^9^ and then comparative studies on all 4 hookworm genomes in 2019 (along with the dog hookworm) and other species of major medical and agricultural importance^10^. Comprehensive stage-specific transcriptional profiles were provided in the genome comparison study^10^ and described in more detail in subsequent studies (e.g. ^11,12^).

However, as with other parasitic nematodes, hookworms rely on multiple organ systems to survive and maintain successful chronic infection within the hosts they infect. Resolution of the cellular diversity and associated molecular differences that underlie functions required for intra-host survival of parasitic nematodes offers insights that may have practical applications, including new targets for preventing transmission and/or controlling infections. The various organ systems that comprise individual parasitic nematodes account for substantial cellular diversity of these pathogens, while the diversity of cellular populations that comprise each organ system remains far less clear. Single-cell expression studies provide a transformative approach to address this knowledge gap. By analyzing the gene expression profiles of individual cells within hookworm organ systems, one could identify distinct cell types and their specific roles in the parasite’s biology. This high-resolution mapping of cellular diversity enables the identification of key molecular pathways and cellular interactions critical for the parasite’s survival and pathogenicity. Nevertheless, there are no single cell transcriptional atlas studies that examine whole organ system heterogeneity, for any parasitic nematode. A recent study provides valuable insights to understand the cellular diversity and molecular variations in *Ascaris suum,* but the study is limited to one specific tissue, the intestine^13^. Another single cell expression study on *Brugia malayi* microfilariae provides the valuable insights into secretory products and anthelminthic responses in different cell types^14^, but this life cycle stage lacks critically important tissue differentiation seen in adults. In the free-living model nematode *C. elegans* Cao *et al* (2017) developed a combinatorial indexing method termed sc-RNAseq, providing significant understanding of L2 stage cellular heterogeneity and development process^15^. The comprehensive single cell atlas of adult *C. elegans* provides a high-resolution map of gene activity, revealing new insights into cell types, gene expression, intercellular interaction and conserved housekeeping gene^16^. These *C. elegans* single-cell studies provide the markers for many cell types of in different nematode species. However, the significant diversification of anatomy and function between *C. elegans* and parasitic nematodes, and consequently, gene and genes families that related to the parasitic lifestyle such as host infection and immune evasion are completely absent or different biological function in *C. elegans*^10,17^ this stipulates more species specific approaches to the study the anatomy and biological process of the host parasitic interaction.

Here, with a goal to reveal the level of heterogeneity at an organ system level, we present the first single-cell transcriptome atlas on adult male and female *Ancylostoma ceylanicum,* a human hookworm. Overall, this study provides a comprehensive cell-type atlas of gene expression in the major tissues of adult hookworms. The results of this study greatly deepens our understanding of the cellular and molecular basis of parasitic nematode survival within hosts. This knowledge can pave the way for the development of innovative strategies to treat, control and prevent hookworm infections, ultimately improving global health outcomes.

## Results and Discussion

### Identification of transcriptionally distinct cell types within male and female hookworm based on multi-omics data integration

Single-cell RNA sequencing was performed on cell suspension of adult female and adult male *A. ceylanicum* worms (collected from Syrian hamsters 20-25 days after oral gavage with infective L3 larvae) using droplet-based 10X Genomics Chromium platform (**Fig 1A**). Transcriptomics data were generated and processed by Cell Ranger (version 7.1.0) from 4,860 and 4,310 adult cells per female and male (respectively), detecting a median of 910 and 378 genes, and 142,868 and 142,448 reads per cell (respectively; **Table 1**).

**Fig 1:**
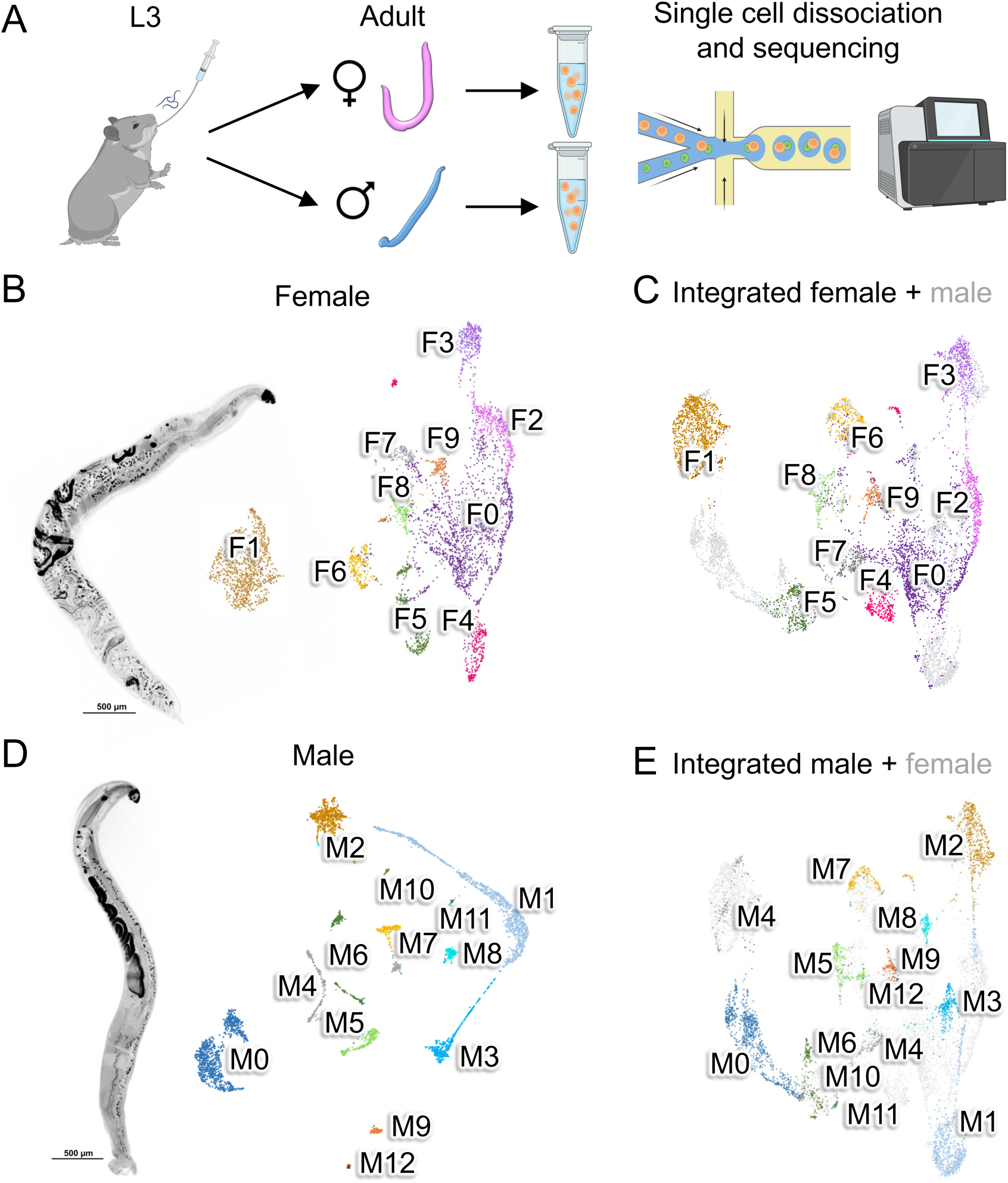
Overview of scRNAseq experiments. (**A**) Schematic of the experimental design. 3-4 week-old Syrian golden hamsters were orally gavaged with 80-100 infective L3 *A. ceylanicum* worms, and adult worms were harvested 20-25 days post-infection. 10 adult female and 10 adult male worms were harvested, and single cell suspensions were prepared from them separately for 10X genomics scRNAseq sequencing. After processing data, UMAP layouts for adult female and male *A. ceylanicum* cells were generated and were clustered using FindClusters, at a resolution of 0.1. Confocal images of adult worms and scRNAseq clustering results are shown for: (**B**) only the female cells; (**C**) the female + the male cells, with female cell cluster identifications according to the female-only analysis in panel A; (**D**) only the male cells; (**E**) the female + the male cells, with male cell cluster identifications according to the male-only analysis in panel D. Clusters are numbered according to the number of cells in each cluster, with numbering starting at 0 and distinct colors assigned per cluster.

**Table 1:** Sample accessions, statistics from Cell Ranger, and statistics after quality control processing.

| Statistic | Adult Female | Adult male |
| --- | --- | --- |
| **Cell Ranger output** |  |  |
| Estimated number of cells | 4,860 | 4,310 |
| Number of reads (million) | 694,337,102 | 613,952,279 |
| Mean reads per cell | 142,868 | 142,448 |
| Median genes per cell | 910 | 378 |
| Fraction of reads in cells | 70.8% | 71.5% |
| Total genes detected | 16,286 | 16,117 |
| Median UMI counts per cell | 2556 | 1,368 |
| **Filtered statistics** |  |  |
| Number of cells | 4,573 | 4,029 |
| Mean UMIs per cell | 2,575 | 1,443 |
| Median genes per cell | 883 | 342 |
| **SRA accession** |  |  |
| BioProject | SRR30221291 | SRR30221289 |
| PRJNA1147629 |  | SRR30221290 |

**Table 2:**
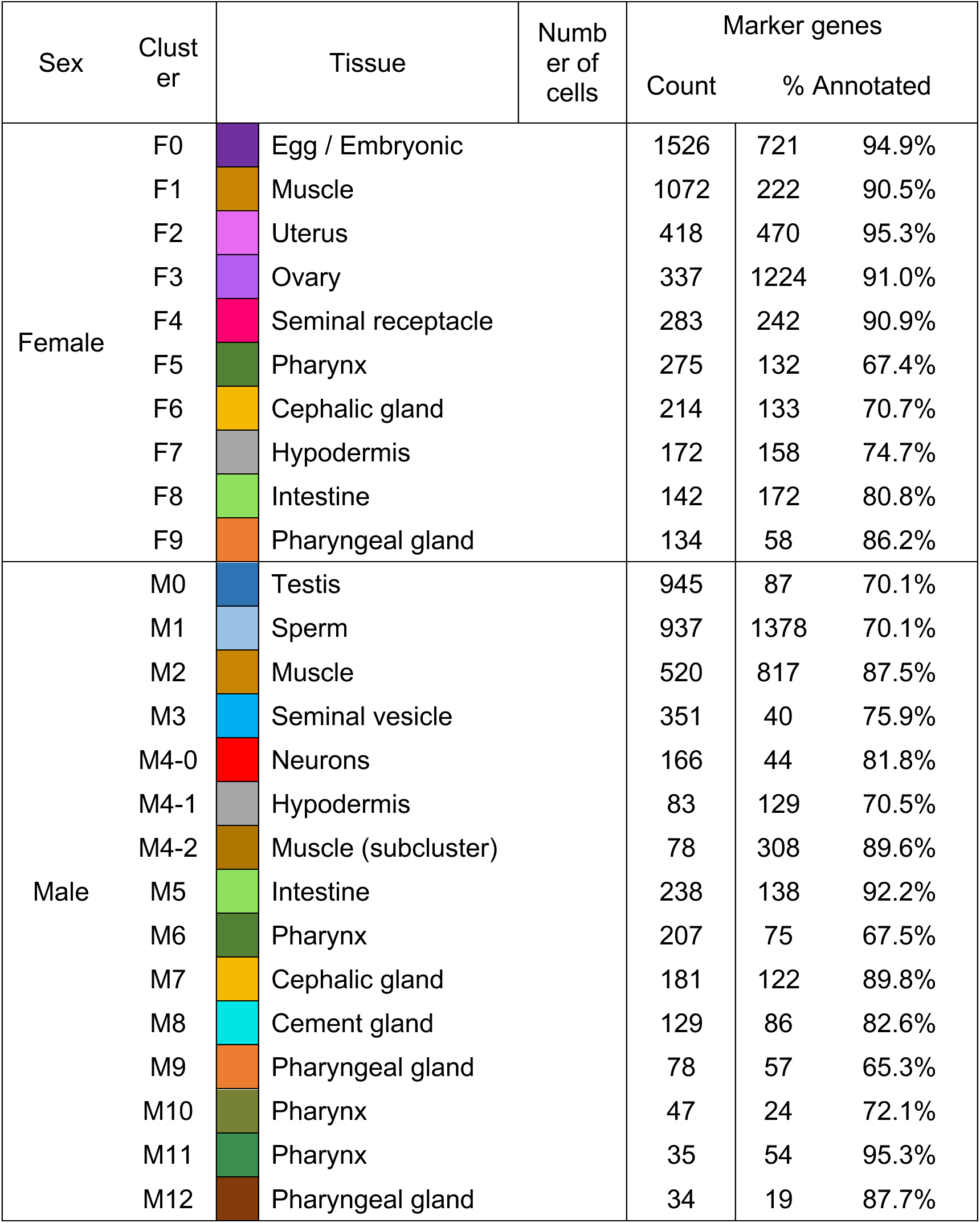
Overview of tissue annotation, cell count and marker gene counts for each cluster.

The cellular maps of the *A. ceylanicum* adults were generated using Seurat^18^, and to better understand cellular diversity and molecular variation in different organ tissues and in different sexes of adult hookworm, we first set out to perform annotation at an organ system level, utilizing all available and informative genomic^10,11^, transcriptomic^10–12,19,20^ and proteomic resources^12^ and robust statistical testing. Adult female cells were clustered into 10 transcriptionally different cell types (F0-F9; **Fig 1B** and **1C**) and adult male cells were clustered into 13 (M0-M12; **Fig 1D** and **1E**), using Seurat resolution value of 0.1, which closely represented the expected number of major tissues while minimizing the possibility of over-resolution. In one instance, a cell cluster with a higher level of apparent heterogeneity was further subclustered with adjust Seurat resolution (M4, as described below).

Final cell type annotations per cluster are shown in **Fig 2**, but the process of annotating them is detailed in the sections below. Integrated clustering with both the male and female cells was performed to help to distinguish non-reproductive and reproductive clusters/tissues for each sex (**Fig 1C** and **1E**; Final tissue annotations in **Fig 3**). We used previously defined marker genes in *Caenorhabditis elegans* to define *A. ceylanicum* marker genes that provided strong information to discriminate between cell populations. We identified and annotated the hookworm marker genes by first identifying reciprocal-best BLAST^21^ orthologs to *C. elegans* (PRJNA13758.WBPS16^22^) and *Ascaris suum*^23^ (PRJNA80881.WBPS17^22^). The identity of *C. elegans* orthologs and their previously-published scRNAseq cell and tissue-specific expression^15^ were used to help assign tissues, in addition to the RNAseq based tissue-specific expression of *A. suum* orthologs (spanning the head, pharynx, intestine and four reproductive tissues^13^) and intestine subcluster marker genes from the *A. suum* intestine scRNAseq study^24^. Marker gene assignment for all genes is provided in in **Supplementary Table S1**, and marker genes separated by cluster and with annotation data and relative abundance statistics are provided in **Supplementary Table S2**.

**Fig 2:**
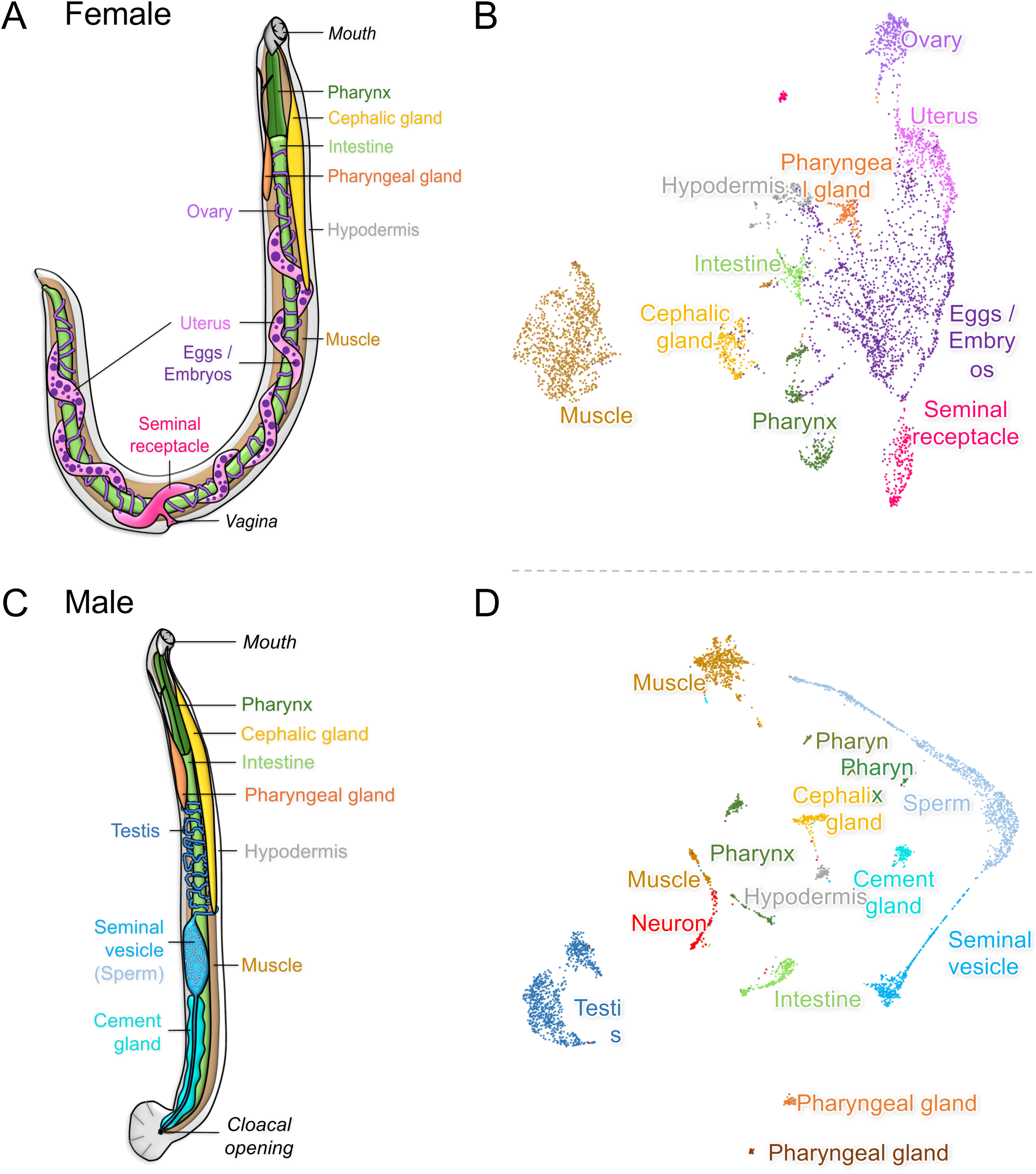
Diagrams of adult *A. ceylanicum* anatomy with the identified tissues labeled, and the final cluster annotations for the sex-specific clustering on UMAP cell layouts. (**A**) The anatomy of the adult female worm; (**B**) The final cluster tissue annotations for the adult female cells; (**C**) The anatomy of the adult male worm; (**D**) The final cluster tissue annotations for the adult male cells.

**Fig 3:**
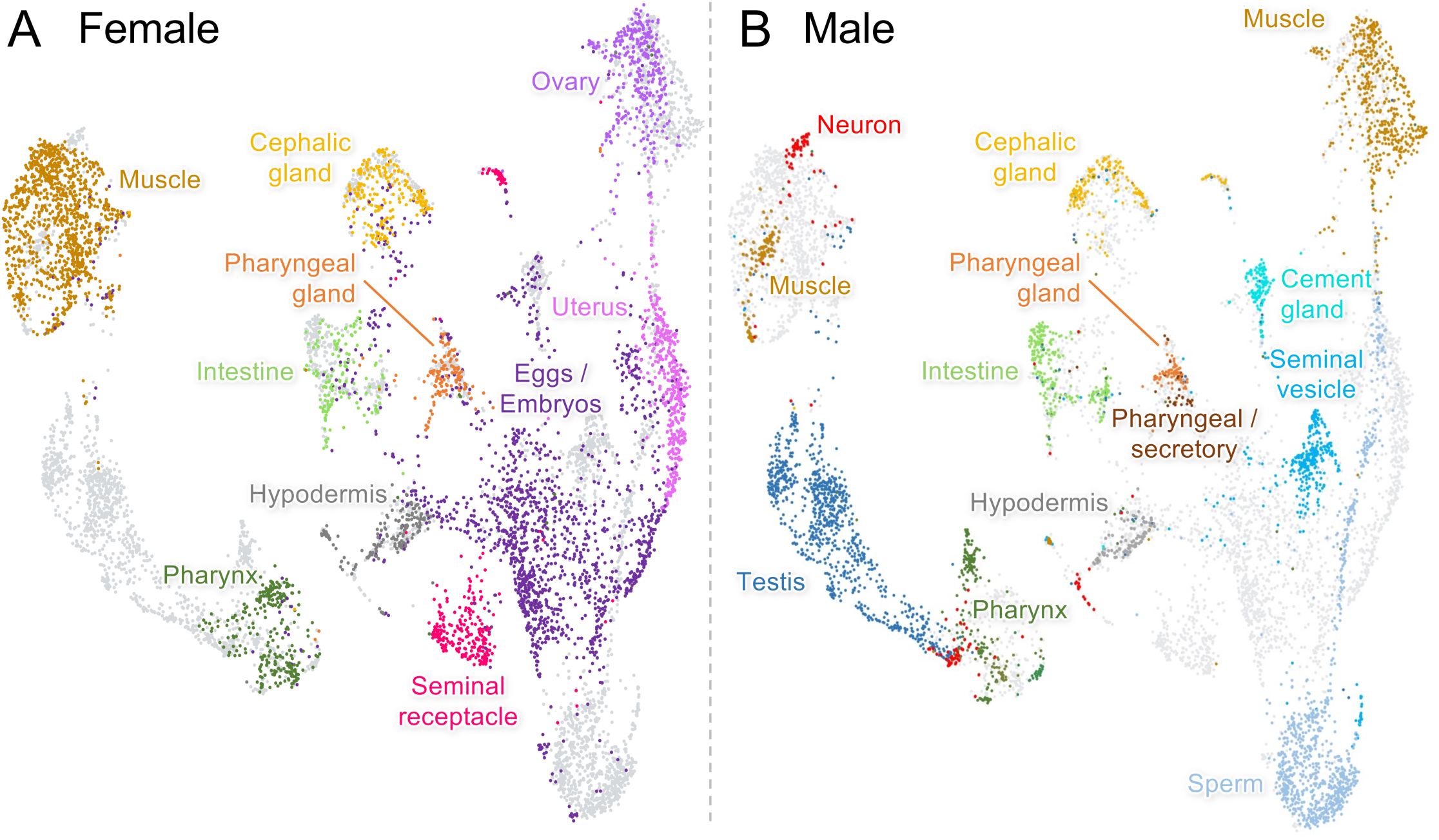
The final cluster annotations overlaid on integrated female + male cell UMAP layouts. (**A**) The final cluster tissue annotations for the adult female cells, overlaid on the integrated female + male cell clustering layout; (**B**) The final cluster tissue annotations for the adult male cells, overlaid on the integrated female + male cell clustering layout.

Marker gene lists per cluster, including the annotations of top markers (ranked by *P* value) and functional enrichment of complete marker gene lists, were used to guide tissue assignment to clusters. These functional annotations for each protein include PANNZER^25^ and Sma3s^26^ protein naming, KEGG^27^ enzyme annotations, InterPro functional domains^28^, gene ontology^29^ assignments, signal peptides for secretion and transmembrane domains^30^. Normalized gene expression levels across the *A. ceylanicum* life cycle^10–12^ and the *A. ceylanicum* male intestine^19^, and adult female and male *A. ceylanicum* ESP proteomics profiles^12^ were also used to guide annotation. Protein conservation data across hookworms and human was also examined using OrthoFinder^31^, as described in our previous study^12^. Complete annotation data is available in **Supplementary Table S1**, significantly enriched KEGG pathways, GO terms and InterPro domains are provided for every marker gene list in **Supplementary Table S2**, and selected significantly enriched GO terms and IPR domains of interest are shown in detail for each annotated tissue in **Fig 4**.

**Fig 4:**
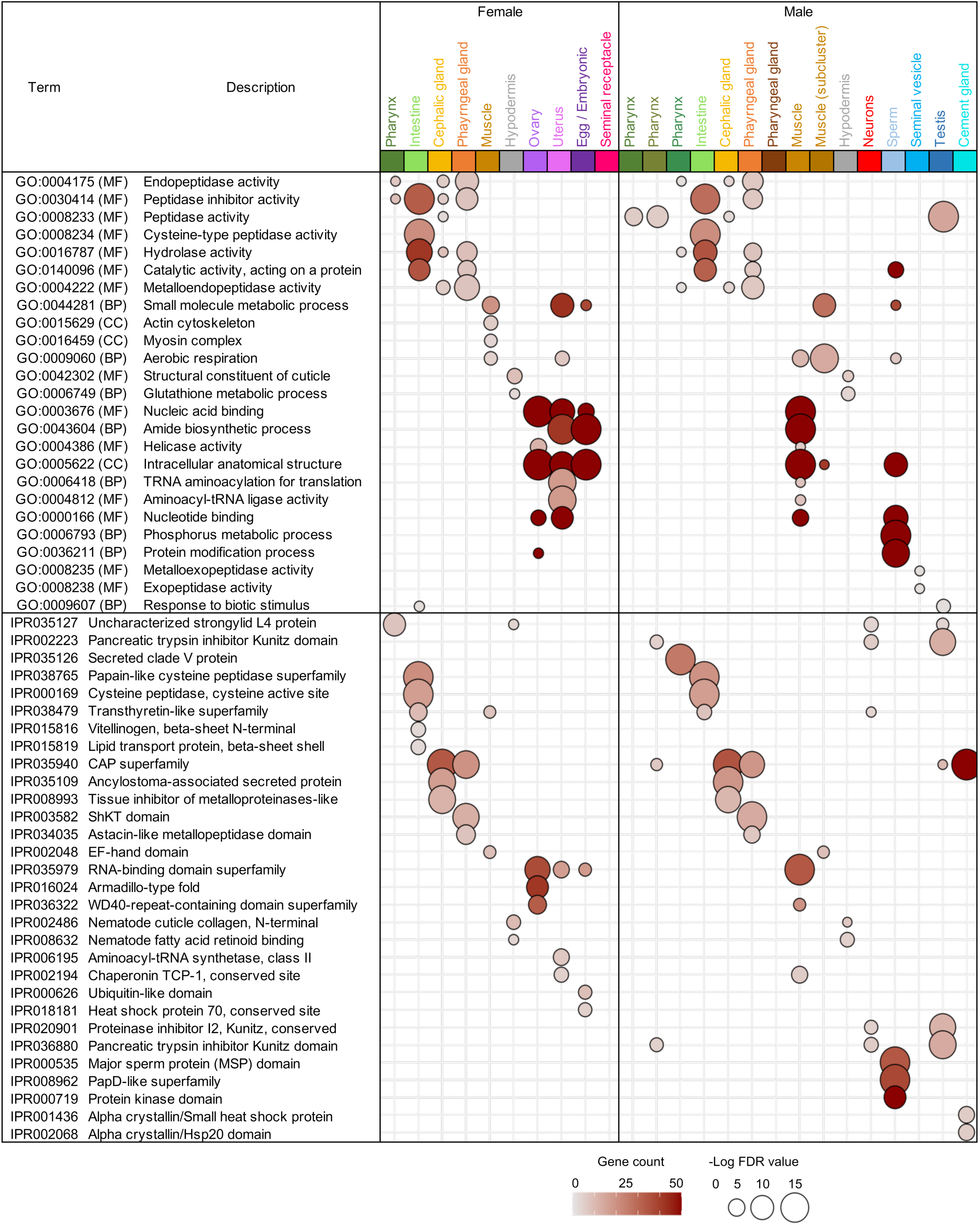
Significance and gene counts for selected enriched Gene Ontology (GO) terms and InterPro (IPR) Domains among each cluster from adult female and male *A. suum.* Terms shown were selected to highlight the important functional annotations for each cluster, and complete enrichment results are provided for each cluster in **Supplementary Table S2**.

### Intestine cells show sex-agnostic patterns of expression

Clusters F8 (142 cells) and M5 (238 cells) overlap in the center of the integrated clustering (**Fig 1C** and **1E**), representing a non-reproductive tissue with shared markers and functional enrichment for intestine. The top shared markers for female and male intestine include several Cathepsin B genes, transthyretins and peptidases (**Supplementary Table S2A**). The top 15 most significant shared marker genes are provided in **Fig 5A** for the intestine clusters, to visualize their relative expression and abundance (and provided as an example, for other tissues). The 172 F8 marker genes were significantly enriched for previous female intestine-overexpressed genes in *A. suum*^13^ (*P* = 4.8×10^-14^) and for intestine tissue marker genes in *C. elegans*^15^ (*P* = 1.8×10^-15^), and the 138 M5 marker genes were significantly enriched for previous male intestine-overexpressed genes in *A. suum* (*P* = 9.9×10^-8^) and for intestine tissue marker genes in *C. elegans* (*P* = 6.8×10^-11^), providing confident evidence in the annotation of these clusters as representing the intestinal tissue. *C. elegans* intestine is composed of a single layer of 20 intestinal cells, and the DNA content of the intestinal nuclei doubles at the end of each larval stage, reaching 32C by the adult stage^32^. We have already demonstrated that the number of intestinal cells in parasitic nematodes is much larger, for example *A. suum* male intestine is comprised of around 1.9 million cells and female intestine around 5.7 million cells^24^. While the number of intestinal cells in hookworms is unknown, our data here indicate that the number is higher than *C. elegans* and there is unrecognized cell heterogeneity and functional diversification between *C. elegans* and among parasitic nematode species.

**Fig 5:**
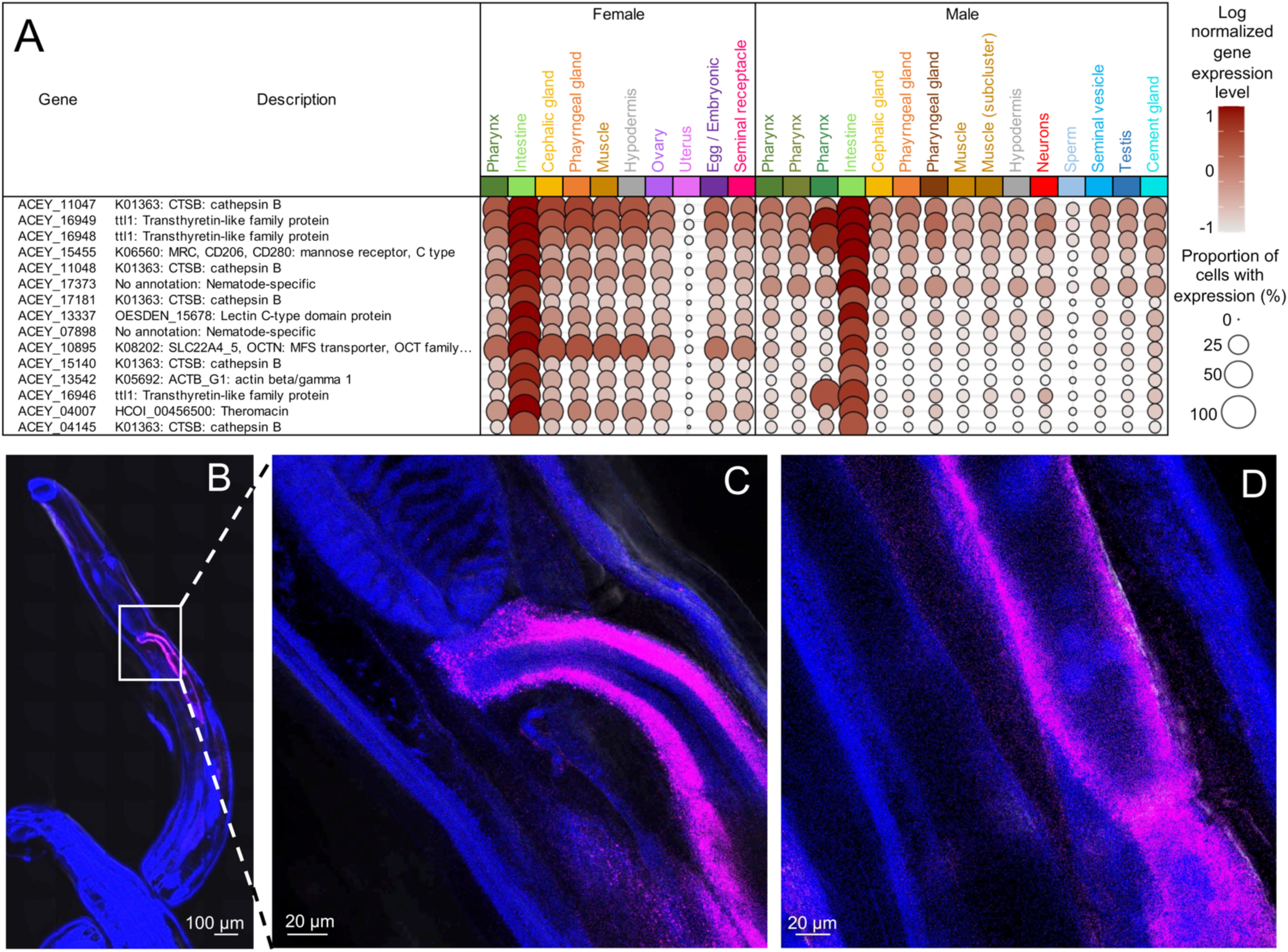
Intestinal cluster markers and validation. (**A**) The average expression level and proportion of cells with expression for the top 15 most significant marker genes for the intestine clusters in both female and male adult *A. suum*. Complete lists of marker genes and their expression data are provided for all clusters in **Supplementary Table S2**. (**B**) Whole-worm fluorescence in situ hybridization (FISH) image showing the expression of *ACEY_10895* (fluorescent signal) throughout the body of the adult hookworm. *ACEY_10895* is stained far red (Alexa fluoro 647), DAPI nuclear stained blue. (**C**&**D**) Magnified view (inset) of the intestine region highlighting strong and specific expression in intestinal cells. The boxed area in panel B corresponds to the region shown in panel C.

Analyzing orthologs from our previous *A. suum* adult female intestinal scRNAseq study^24^, we identify significant enrichment for markers of all three major intestinal clusters (As-C1, C2 and C3) among the intestinal marker genes from both sexes, with the strongest enrichment (*P* = 9.2×10^-7^ for females and *P* = 2.4×10^-10^ for males) for the As-C3, consistent with this cluster’s high conservation across the phylum Nematoda and core intestinal functions such as peptidase and metabolic activity, and the lowest enrichment for As-C2 (*P* = 1.1×10^-3^ for females and *P* = 2.3×10^-2^ for males), consistent with the clade III-specificity of its marker genes (hookworms are clade V compared to *A. suum* taxonomically positioned in clade III^10^), with functions including calcium signaling and transcription regulation^24^. The top As-C3 marker, *vit-6*, was the third-ranked marker in the female *A. ceylanicum* intestine but was not a significant marker in the male intestine and was (i) ∼10,000-fold higher-expressed in the whole-worm adult female *A. ceylanicum* vs male^10,11^, (ii) ∼389-fold higher-expressed in the *A. suum* female intestine vs the male^13^ and (iii) 2.6-fold higher expressed in the *Toxocara canis* female intestine vs the male^33^, suggesting female-specific intestinal expression to a degree that varies by nematode species. This gene has potential roles in hormone signaling, innate immune responses and pathogen recognition but its function in the nematode intestine is not well understood despite its consistently high detection in several species^33^.

Both male and female marker genes for F8 and M5 were strongly significantly enriched for cysteine peptidases (*P* < 10^-15^), proteolysis (*P* = 1.2×10^-13^ and *P* = 3.0×10^-14^, respectively), and hydrolase activity (8.7×10^-12^ and 6.7×10^-10^), as well as many other intestine-associated protein domains, KEGG pathways and Gene Ontology terms (**Fig 4**; **Supplementary Table S2A**). The intestine cell type showed strong enrichment for various proteolytic enzymes, including cathepsin B, cathepsin A, cytosolic nonspecific dipeptidases, metallopeptidases, and peptidase family enzymes. This enrichment aligns with the established roles of these proteases in intestinal digestion and nutrient acquisition. In hematophagous nematodes such as *Ancylostoma* and *Haemonchus contortus*, digestive proteases like cathepsins, metallopeptidases, and aspartic proteases (e.g., APR-1) operate in a coordinated cascade for hemoglobin degradation. These enzymes facilitate nutrient acquisition from host blood meals. Specifically, cathepsin B, a cysteine protease, is secreted by adult hookworms and localized to the intestinal brush border in *Ancylostoma caninum*. It plays a critical role in cleaving and denaturing native hemoglobin^34^, and cathepsin B cysteine proteases are promising vaccine targets against blood-feeding gastrointestinal nematodes, including hookworms, due to their essential role in parasite blood digestion^35^. *AceyCP1* (intestinal cathepsin B protease) was shown to provide significant protection in vaccination trials using recombinant protein in Syrian hamsters, reducing worm burden by ∼50–65% and impairing hookworm motility by neutralizing blood digestion within the intestinal lumen^35^. Here, 14 of the top 50 shared female and male intestine markers (including the top shared marker *ACEY_11047*) were cathepsin B proteins with no *C. elegans* orthologs, providing specific targets for future study of these critical hookworm proteins.

To validate the expression of gene enriched the intestine cluster identified in the scRNA seq data, we performed a FISH assay to validate the tissue-specific expression of a key marker gene, against a background of DAPI staining for the adult worm nuclei. We targeted the intestine for this analysis since it has a well-defined anatomy (**Fig 6A**) and is in direct contact with the outside environment. Among the top intestinal marker genes, we prioritized *ACEY_10895,* the top most significant marker in the females (29^th^ in the males), with the highest average expression among the shared female and male intestinal markers, and 17.59-fold and 6.56-higher expression in the female and male intestine vs other tissues, respectively. *ACEY_10895* is an organic cation transporter family members 4/5 (OCTN1/OCTN2), belonging to the solute carrier family 22 (SLC22) and classified as a major facilitator superfamily (MFS) transporter^36^. Strong FISH signal was observed in the intestinal region of adult hookworms (**Fig 5B-D**), with strong and specific fluorescence localized to the intestinal cell wall, consistent with scRNAseq transcriptional distinct cell type. Negative controls (including no-RNA probes and no probes/no hair pins) showed no fluorescence signal, confirming the specificity of the FISH assay. These results validate the scRNAseq expression, demonstrating that the predicted enrichment of marker gene corresponds to their spatial localization in the intestine of adult hookworms.

**Fig 6:**
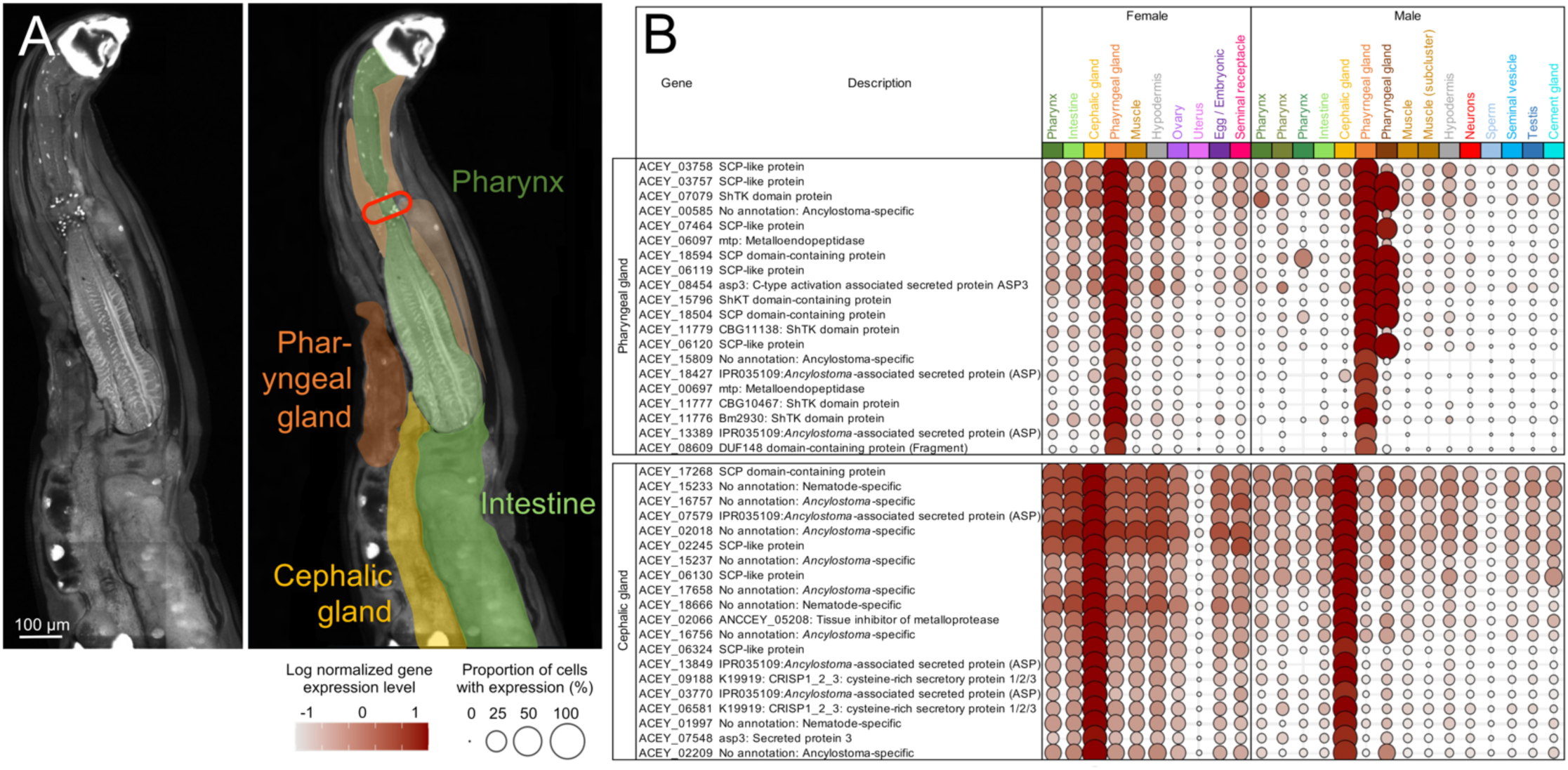
Secretory gland identification and cluster marker genes. (**A**) Confocal microscopy showing the distinct non-reproductive tissues in the anterior end of an adult female *A. ceylanicum* worm. Color overlays on the right panel indicate tissues corresponding to tissues identified in the scRNAseq clustering analysis. Red line = nerve ring. (**B**) The average expression level and proportion of cells with expression for the top 20 most significant marker genes for the pharyngeal gland and cephalic gland clusters in both female and male adult *A. suum*. Complete lists of marker genes and their expression data are provided for all clusters in **Supplementary Table S2**.

### Identification of three distinct populations of pharyngeal cells in a male hookworm

In close proximity to the intestine clusters (**Fig 1**) were F5 (275 cells) and M6 (207 cells), M10 (47 cells) and M11 (35 cells). The 132 markers for F5 were significantly enriched for pharynx tissue (*P* = 9.9×10^-8^) and cells (*P* = 3.0×10^-4^) based on *C. elegans* orthologs^15^ as well as female ESPs^12^ (*P* = 5.1×10^-7^; **Supplementary Table S2B**). Clusters M6, M10 and M11 were all enriched for male ESPs (*P* = 4.8×10^-4^, *P* = 3.5×10^-4^ and *P* = 7.6×10^-8^, respectively), with cluster M6 (75 markers) being enriched for functions including C-type lectins (*P* = 6.1×10^-9^) and peptidase inhibitor activity (*P* = 9.8×10^-6^), cluster M10 (24 markers) being enriched for endopeptidase inhibitor activity (*P* = 4.6×10^-8^) and cluster M11 (54 markers) being enriched for Secreted clade V proteins (*P* < 10^-15^; **Supplementary Table S2B**). The division of this apparent pharynx cells cluster into three subclusters in the male may reflect the identification of distinct pharynx cell types that were not well-separated transcriptionally in the F5. The integrated clustering layout reflects this, showing the female pharynx cluster being spread out across the region between the three male clusters (**Fig. 3A** and **3B**). We were concerned that splitting this apparent male pharynx cluster into three clusters was reducing the significance of pharynx marker genes in each cluster; i.e., if a pharynx marker gene was highly expressed in all three then it won’t be a significant marker in any of them. Therefore, to make a comparison to the female pharynx cluster, we combined the cells belonging to M6, M10 and M11 into a single pharynx cluster representing 289 cells. When these clusters are merged, we identify 232 significant marker genes and we identify significant enrichment for pharynx tissue (*P* = 0.046) and pharyngeal gland cells (*P* = 0.022) based on *C. elegans* orthologs^15^, and we still see significant enrichment of male *A. ceylanicum* ESPs^12^ (*P* < 10^-15^), including ESPs that showed no gene expression in the adult male intestine^19^ (*P* = 3.6×10^-5^).

As seen in the *A. suum* pharynx^13^, both the male and the female pharynx clusters were enriched for many peptidase-related functional enrichment terms, including the general terms “endopeptidase activity” and “proteolysis” in the single female cluster F5, “metallopeptidase activity” in M11, and peptidase inhibitor activity in M6 and M10. The marker gene *nas-5* (*ACEY_06769*) was the top shared pharynx marker, being the 6^th^ highest marker gene in F5 and the 21^st^ highest marker gene in the male merged pharynx cluster (**Supplementary Table S2B**), and it has been previously identified in the pharyngeal glands of *C. elegans*^15^. Among the top shared pharynx markers, we also identify three *transthyretin-like* proteins: *ttr-1* / *ACEY_18247* (ranked 2^nd^), *ttr-2* / *ACEY_09447* (ranked 3^rd^) and *Transthyretin-like family protein* / *ACEY_02199* (ranked 6^th^). Transthyretin-like proteins are one of the largest conserved nematode-specific gene families^37^, they were the only protein family to be detected in the ESPs across 20 distinct nematode species^38^, and were significantly enriched among species-specific intestine-expressed protein families^20^ but their roles are poorly understood. Here, we identify three with consistent pharynx-specific expression in both females and males.

Overall, we identify sex-specific differences in the pharynx cells, including distinct transcriptional subdivision of pharynx tissue in the adult male relative to the female, with overall well-supported annotations and shared pharynx-specific marker genes of interest.

### Pharyngeal and cephalic secretory gland cells are distinct, highly hookworm-specific, and conserved across sexes

Using confocal microscopy, we identified distinct cephalic and pharyngeal glands at the anterior end of adult worms (**Fig 6A**), aligning with previously described *A. ceylanicum* anatomy, where the pharyngeal gland was referred to as an “excretory gland” ^39^. The cephalic gland has a substantially larger anatomical size relative to the pharyngeal gland, extending far beyond the beginning of the intestine, as previously described^39^, and was therefore expected to be represented by clusters with more cells than the pharyngeal gland. The prominent pharynx was observed as an elongated structure at the anterior region with a clear point of attachment to the intestine, with the pharyngeal gland being positioned entirely beside the pharynx.

There was 90% overlap between the marker genes from F9 (134 cells, 58 markers) and M9 (78 cells, 57 markers; **Supplementary Table S2C**). These marker genes were not enriched for any tissues or cell types from the *C. elegans* dataset^15^, with only one gene (*ACEY_06281*) having a *C. elegans* ortholog, which was associated with excretory glands (*W02D7.3, smex-1; Small Membrane protein expressed in EXcretory gland 1*). Among the 52 marker genes overlapping in both F9 and M9, only 8 had reciprocal orthologs in the hookworm *N. americanus* and 30 (57.7%) had reciprocal orthologs in *A. caninum*, suggesting rapid evolution and species-specificity.

Marker sets from both of these clusters were significantly enriched for *A. ceylanicum* ESPs^12^ which showed no gene expression in the adult intestine^11,19^ (*P* < 10^-15^), accounting for 29.3% and 31.6% of marker genes, respectively (compared to a maximum of 0.66% across non-secretory tissues). This evidence suggests that this cell type represents a secretory gland. Among the top markers in these clusters was *A. ceylanicum* anticoagulant protein 1 (AceAP-1, *ACEY_09599*), which is the closest ortholog in *A. ceylanicum* to *A. caninum* Ac-AP-12, identified only in the adult *A. caninum* pharyngeal glands^40^, providing support that these were pharyngeal gland cells. Hookworm have developed potent mechanism to disturb host hemostatic mechanism and facilitate the blood feeding. Recombinant Ac-AP12 showed anticoagulant activity in human blood plasma by specifically inhibiting the activity of human factor Xa^40^. In a previous proteomics-based identification of immunoreactive proteins from the pharyngeal gland in *Anisakis simplex*^41^, many SCP/CAP-domain proteins and metalloendopeptidase proteins were identified, and we also see the most significant functional enrichment for both of these protein families in both male and female marker gene lists (*P* ≤ 6.3×10^-9^ for all comparisons). The smaller cluster M12 (34 cells, 19 markers) closely positioned to M9, was also enriched for ESPs with no intestine expression^19^ (*P* = 5.0×10^-7^), and its most significant marker of 19 markers (**Supplementary Table S2C**), was also a marker in F9 (*ACEY_08986*, putative Beta-lactamase). Because of this evidence, M12 was annotated as a pharyngeal gland tissue, representing either a distinct gland or distinct cell types within a single gland.

F6 (214 cells, 133 markers) and M7 (181 cells, 122 markers) represents a larger secretory gland cluster, with no significant enrichment with marker genes for cells or tissues from *C. elegans*^15^, and strong enrichment for detected *A. ceylanicum* ESPs (76% of female markers, and 80.3% of male markers; *P* < 10^-15^ for both; **Supplementary Table S2D**). In *A. caninum*, ASP-6 localized to the cephalic gland^42^, and in our *A. ceylanicum* genome annotation, we identify 15 ASP-6 orthologs, two of which (*ACEY_07543* and *ACEY_07544*) were shared female and male markers (**Supplementary Table S2D**) in clusters F6 and M7. No markers in F9 and M9 clusters (the pharyngeal gland cluster, described above) were ASP-6 orthologs. This provides evidence that F6 and M7 represent the cephalic gland.

The pharyngeal and cephalic gland cell clusters were substantially overlapping between male and female, and were positioned distinctly from other tissues in both sexes (**Figs 3A** and **3B**), but both pharyngeal and cephalic gland marker cells in both males and females were strongly enriched for secreted CAP / SCP / venom allergen-like domain proteins in nematodes, which are thought to be involved in a wide range of critical host-survival processes including modulation of the host immune system^43^, and were the most strongly enriched protein family in adult *A. ceylanicum* ESPs^12^. Here, for the first time we characterize highly abundant but distinct CAP domain gene expression from each of the pharyngeal and cephalic glands (top 15 marker genes for each cluster shown in **Fig 6B**).

Among marker genes for these cell clusters, the cephalic gland but not the pharyngeal gland in both female and male were significantly enriched for functions including *Ancylostoma*-associated secreted proteins (ASPs; IPR035109; previously among the most significantly enriched protein families in *A. ceylanicum* adult ESPs^12^) and tissue inhibitor of metalloproteinases (TIMPs; IPR008993; a previously described TIMP was among the most abundant proteins released from adult *A. caninum*^44^; **Fig 4**). In a mouse model of ancylostomiasis, immunization with recombinant ASP-1 significantly reduced worm burden in the lungs by 79% when alum was used as an adjuvant^45^, highlighting the utility of this protein family in infection prevention. Here, we provide the first evidence that these protein family members are primarily secreted specifically from the cephalic gland.

Meanwhile, the pharyngeal gland but not the cephalic gland in both female and male was enriched for ShKT domain proteins (IPR003582; the most abundant protein domain identified across parasitic nematode ESPs^46^, with possible roles in host-parasite interactions^47^) and metallopeptidases including Astacin-like metallopeptidases (IPR034035; responsible for a wide range of functions including tissue migration^48^; **Fig 4**). Three additional *A. ceylanicum* anticoagulant proteins (AceAP-1, peptides 1, 4 and 12) were shared female and male markers in the pharyngeal gland but not the cephalic gland, highlighting a potential role in blood feeding for this gland relative to the cephalic. These protein families have been previously associated specifically with the pharyngeal gland in any previous research.

Overall, the results here provide that both shared and specific gland-associated genes and gene families are present, providing empirical support at a resolution which has not been previously reported. These results may offer specific novel insights into the functions provided by these cells to complement the present very broad functional understanding.

### Identification of other non-reproductive tissues, the hypodermis and body wall muscle

M4 (327 cells, 481 markers) and F7 (172 cells, 158 markers) were mostly overlapping in the integrated clustering (**Fig 1C** and **1E**), with small number of cells from the M4 found overlapping the F1 (1072 cells, 222 markers) and the pharynx clusters (F6 and F9) described above. Because of this spread of the cells in the male, we performed subclustering of M4 cells using a resolution of 0.2, to facilitate distinction among cells and functions enriched in these three distinct subclusters (**Fig S1A**), which were identified in distinct regions of the male-only clustering (**Fig S1B**). The largest subcluster M4-0 (166 cells, 44 markers) was significantly enriched for *C. elegans* markers of flp-1(+)_interneuron cells (*P* = 4.8×10^-4^), and its 44 markers included two annotated neuron-associated genes: PDF-1 (**Supplementary Table S2E**), which functions in non–sex-specific neurons to produce male-specific, goal-oriented exploratory behavior^49^, and *sul-2*, which is only expressed in sensory neurons and is associated with longevity^50^. The male neuron cells from M4-0 were found across different major clusters in the integrated clustering (**Fig S1C**), possibly indicating tissue-specific neurons which fail to be resolved in the female sample (which tended to have more diffuse clusters overall). Subcluster M4-1 (83 cells, 129 markers; **Fig S1C**) was significantly enriched for orthologs of *C. elegans* markers^15^ of glia (*P* = 3.5×10^-3^) and hypodermis (*P* = 7.4×10^-3^) and overlapped F7 (**Fig S1C**) which was also significantly enriched for hypodermis tissue (172 cells and 158 markers, **Fig S1A**) based on *C. elegans* markers^15^ (*P* = 2.7×10^-3^). The most significant marker gene (*ACEY_02130*) in both M4-1 and F7 was OV-17 (**Supplementary Table S2F**), which localizes to the hypodermis of adult *Onchocerca volvulus*^51^. These two clusters overlap on the integrated clustering (**Fig S1C**) and share 90 gene markers (57.0% and 69.8% of female and male markers, respectively), so these two subclusters were annotated as hypodermis tissue.

F7 did not contain cells that were spread out across the female-only clustering, with all of the cells positioned close to clusters F8 (intestine) and F9 (pharynx) in the UMAP integrated clustering (**Fig 2A**). To test whether we would observe similar results to the subclustering of M4, we also performed subclustering of F7 at the same resolution (0.2), resulting in identification of two subclusters. However, unlike the male, there was no clear muscle-annotated subcluster, with the larger (102 cells and 63 marker genes) of the two subclusters having marker genes and functional enrichment overlapping the overall F7, and the smaller (76 cells) having no significant enrichment and no functional enrichment among the 6 marker genes (data not shown). We therefore observed no statistical justification for sub-clustering the female hypodermis cluster, as we performed for the male.

Muscle tissue cells were annotated across different clusters in male and female worms. In adult female worms, in addition to body wall muscle tissue, there are smooth muscle-like myoepithelial sheath cells in the proximal ovary^52^ (F3, described below), and the uterus is also surrounded by sex-specific muscles that contract in order to facilitate egg laying^53^ (F2, described below). However, the transcriptionally distinct F1 (1072 cells, 222 markers) was confidently annotated as muscle tissue based on: (i) marker genes overlapping with markers for body wall muscle cells and tissues from *C. elegans*^15^ (*P* < 1×10^-15^ for both), (ii) the functional enrichment of many muscle-related terms including “actin cytoskeleton” (GO:0015629; *P* = 8.7×10^-4^) and “myosin complex” (GO:0016459; *P* = 3.3×10^-3^) and (iii) specific muscle-associated marker genes including troponin T, troponin C and troponin I **Supplementary Table S2G**), the three components of the broadly-conserved troponin complex that is the major regulator of muscle contraction^54^, and (iv) the marker gene paramyosin, a major structural constituent of nematode body wall muscle^55^. Subcluster M4-2 (as described above) overlapped with this female muscle cluster (F1) in the integrated clustering, and shared 48.2% of the female muscle cluster’s marker genes including 31 that are orthologs of genes most highly expressed in *C. elegans* body wall muscle, and it was significantly enriched for orthologs of *C. elegans* markers of muscle cells and tissue^15^ (*P* < 1×10^-15^ for both) **Supplementary Table S2G**). The conservation of this integrated cluster in both males and females, as well as the marker genes identified, support that M4-2 represents non-reproductive body wall muscle tissue.

The larger M2 (520 cells, 817 markers) clusters distinctly far away from the male subcluster M4-1, but also shows significant enrichment for orthologs of markers of muscle tissue and cells from *C. elegans*^15^ (*P* = 9.3×10^-7^ and 5.4×10^-4^, respectively; **Supplementary Table S2G**), as well as for sex myoblast cells and gonad tissue (*P* < 1×10^-15^ for both). These results support that M2 is male reproductive-associated muscle tissue, further supported by the cluster’s overlap with ovary tissue from the females (F3, described in detail below), which contains muscle-like myoepithelial sheath cells^52^. Based on these observations, M2, which was positioned at the top-right of the integrated clustering, likely represents reproductive tissue-associated muscle cells with transcriptional profiles distinct from body wall muscle cells (M4-2) that were positioned at the top-left of the integrated clustering.

### Transcriptionally distinct female reproductive tract tissues and associations with muscle tissue

For the annotation of reproductive tissues, known *C. elegans* marker genes were used to help guide annotation as for the other tissues. However, it is important to note that *C. elegans* is hermaphroditic^56^, which introduces fundamental differences in reproductive biology compared to hookworm. *C. elegans* marker genes were annotated as being tissue-specific, and not sex-specific, for this reason. Cells from cluster F2 (418 cells, 470 markers; **Fig 1B**) were significantly enriched for *C. elegans* markers of gonad tissue^15^ (*P* < 1×10^-15^) and for orthologs of *A. suum* uterus-overexpressed genes^13^ (*P* = 3.1×10^-3^; **Supplementary Table S2H**), supporting F2 annotation as uterus cells. The nematode uterine wall is thick and muscular^57,58^, so it is not surprising that marker genes for F2 were also significantly enriched for *C. elegans* markers of body wall muscle^15^ (*P* < 1.6 ×10^-8^). In that dataset, no distinction was made between reproductive-associated muscle cells and body wall muscle cells. Significant functional enrichment for F2 includes 11 GO terms that were also highly significantly enriched in uterus tissue in *A. suum*^13^, including (i) nucleotide binding (GO:0000166; *P* = 1.0×10^-4^, 2nd ranked in the previous study; *P* = 1.3×10^-8^ in the current study), (ii) tRNA aminoacylation for protein translation (GO:0006418; *P* = 1.1×10^-4^, 3rd rank in the previous study; *P* = 2.6×10^-13^ in the current study), and (iii) aminoacyl-tRNA ligase activity (GO:0004812; *P* = 1.2×10^-4^, 4th rank in the previous study; *P* = 2.5×10^-13^ in the current study). Taken together, these results strongly suggest that F2 represents the *A. ceylanicum* uterus.

Cells from cluster F3 (337 cells, 1224 markers; **Fig 1B**) were significantly enriched for *C. elegans* markers of gonad tissue^15^ (*P* < 1×10^-15^) and germline tissue (*P* < 1×10^-15^), and for orthologs of *A. suum* ovary-overexpressed genes^13^ (*P* = 3.9×10^-9^; **Supplementary Table SI**), providing evidence that F3 cell cluster represent the ovary. Significant functional enrichment for F3 includes 11 GO terms that were also highly significantly enriched in uterus tissue in *A. suum*^13^. These included 8 of the top 10 terms from the previous study, including four terms related to binding, helicase activity (GO:0004386; *P* = 5.5×10^-6^, 7th rank in the previous study; *P* = 4.9×10^-5^ in the current study) and intracellular anatomical structure (GO:0005622; *P* = 6.1×10^-5^, 8th rank in the previous study; *P* = 6.1×10^-20^ in the current study). F3 overlapped with the primary male muscle cluster (M2) on the integrated clustering (**Fig 3B**), but the enrichment analysis does support its identification as the ovary. As described in the above muscle tissue section, the male muscle cluster may therefore also represent reproduction-associated muscle cells.

F4 (283 cells, 242 markers) represents a female-specific cluster (**Fig 1C** and **1E**) which was most significantly enriched for *C. elegans* markers of gonad tissue^15^ (*P* < 1×10^-15^; **Supplementary Table S2J**), indicating that it was a female reproductive tissue represented by fewer cells than the ovary or uterus, but more cells than other tissues including the pharynx and intestine. Based on the expected major female reproductive tissues in *A. ceylanicum*^59^, we have annotated F4 as the seminal receptacle, the widening at the beginning of the uterus storing sperm cells but not the site of embryogenesis^60^ (**Fig 7B**). The only other significant enrichment for this tissue was for *C. elegans* markers of oxygen sensory neurons^15^, but the large number of cells and their lack of overlap with male cells in the clustering makes it unlikely that F4 represented only neurons. The lack of known seminal receptacle-specific marker genes makes it difficult to further support this annotation, but provides the first such list of marker genes, and includes histones H3.3 and his1, as well as histone chaperone protein asf1.

**Fig 7:**
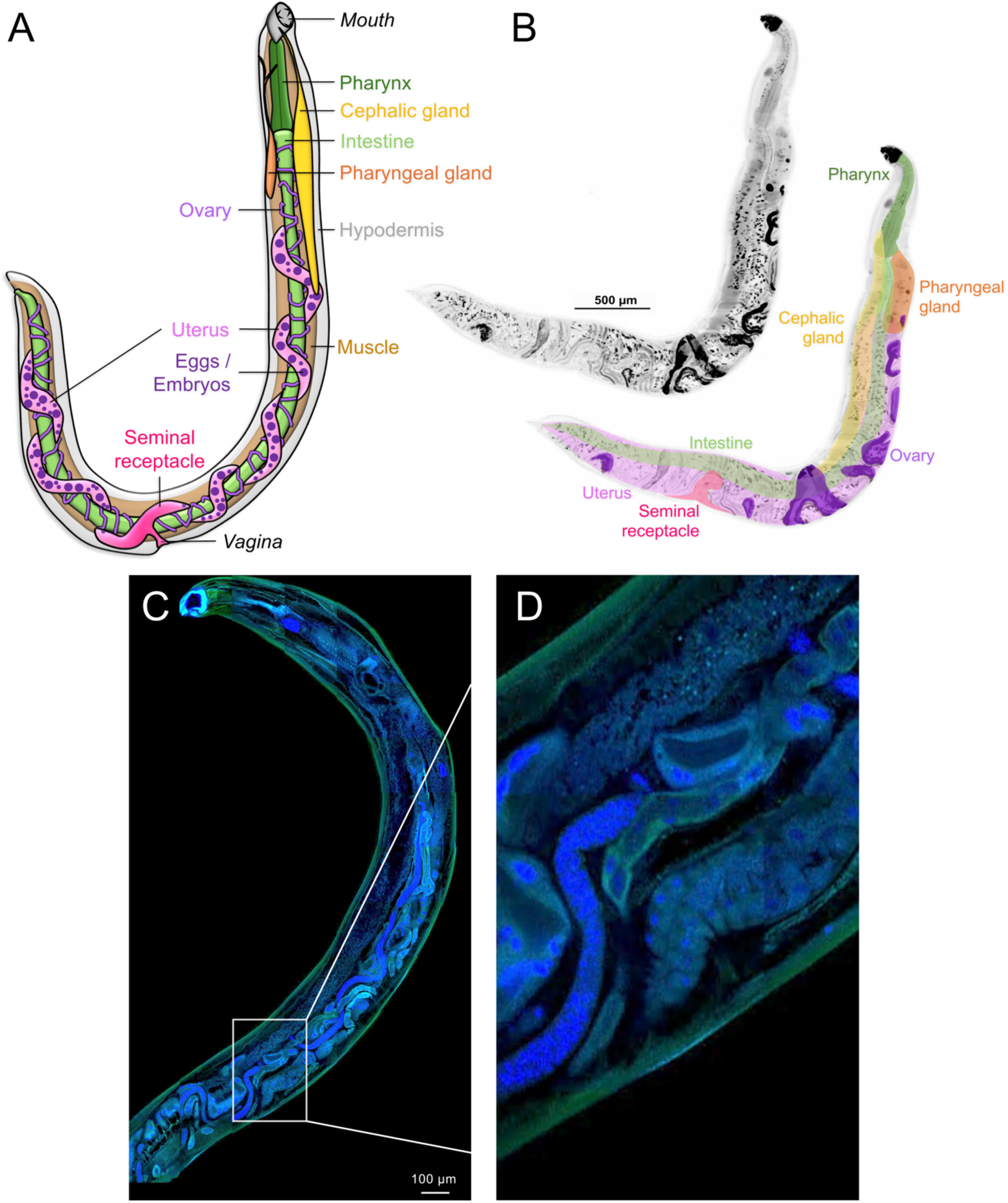
Microscopy of adult female *A. ceylanicum* worms indicating major tissues of interest. (**A**) Schematic representation of a female adult worms with major tissues indicated. (**B**) Confocal image of an adult female *A. ceylanicum* worm stained with BB, acquired at 20x magnification showing distinct anatomical features include the pharynx, pharyngeal gland, cephalic gland, intestine and reproductive organs including the ovary, seminal receptacle and uterus. (**C**) Confocal microscopy image of an adult female *A. ceylanicum* worm stained with DAPI and phalloidin revealing detailed visualization of cytoskeleton structure, including the actin filaments of the ovary and uterus. (**D**) Increased magnification of a region of the adult female worm indicated with a white box on panel C, showing evidence of eggs (bright blue) in the ovary and uterus.

Cells from cluster F0 (1526 cells, 721 markers; **Fig 1A**) were widely spread out across the cluster space, indicating a variety of transcriptional profiles in these cells. F0 was enriched for *C. elegans* markers^15^ of vulval precursors (*P* = 6.0×10^-15^) and myoblasts (*P* = 1.1×10^-10^), indicating that these cells represent early developmental stages, indicative of embryos in different developmental stages. The third most significant marker gene for F0 was an ortholog of *C. elegans CBD-1*, which is required for eggshell vitelline layer formation and egg activation^61^, and the 2^nd^, 7^th^ and 8^th^-ranked markers were chondroitin proteoglycan proteins (**Supplementary Table S2K**), which are essential for embryonic cell division in *C. elegans*^62^. Additionally, the top functional enrichment for F0 includes many terms associated with rapid protein production and biosynthesis including the KEGG pathway for Ribosomes (*P* < 1×10^-15^) and many strongly enriched (*P* < 1×10^-15^) GO terms associated with biosynthetic processes. Taken together, all of this evidence indicates that F0 represents cells from developing eggs and embryos within the adult female.

**Fig 7** shows a schematic representation (**Fig 7A**) of all the described female tissues, supported by structural observation of BB-stained female hookworm using confocal microscopy. While BB staining effectively highlighted the nuclei within the reproductive tissues, the ovary and uterus could not be clearly distinguished solely based on BB staining due to close anatomical association and lack of distinct morphological features using BB staining. Counterstaining with phalloidin provided detailed visualization of the cytoskeletal structure within the hookworm including the ovary and its surrounding tissues. The phalloidin highlighted the actin filaments forming the structural framework of the ovary and uterus, allowing for precise delineation of these tissues (**Fig 7C** and **7D**) further supporting these observations, demonstrating apparent staining of developing eggs throughout the ovary and larger eggs / embryos in the uterus. The female reproductive tract, based on the confocal images showed the ovary as a long thin tube-like structure coiled over the intestine, that continues with tubular uterus and seminal receptacles. Seminal receptacles appeared as a sac-like structure at beginning of ovary.

### The male reproductive tissues are defined by transcriptionally distinct clusters representing sperm, testis, seminal vesicle and cement gland clusters

M1 (937 cells, 1378 markers) contained the highest number of markers than all other cell clusters (1,378 genes), indicating that it was a very distinct cell type. The top 5 markers were all “major sperm protein” (MSP) genes, and the “MSP domain” (IPR000535) and the related “PapD-like superfamily” (IPR008962) InterPro domains were the most significantly enriched of all domains, pathways and terms among these marker genes (*P* < 10^-15^ for both; **Supplementary Table S2L**). Major sperm proteins comprise the filaments of pseudopod of sperm and are assembled during spermatogenesis^63^. Top markers for M1 also include an ortholog of *alg-3* (6^th^-ranked), which is critical for sperm development and thermotolerance^64^, and an ortholog of *spe-4* (28^th^-ranked), which encodes transmembrane protein localized within vesicles of spermatids and spermatozoa^65^. Markers from this cell cluster were enriched for gonad marker genes in *C. elegans* orthologs^15^ (*P* = 5.5×10^-7^), as well as overexpression in the testis in *A. suum* orthologs^13^ (*P* < 10^-15^). This finding from *A. suum* is thought to result from the fact that sperm and testis were not separated in that previous study, so gene overexpression profiles assigned to the “testis” tissue may better represent sperm transcription. Nonetheless, the marker gene lists and functional enrichment overwhelmingly support M1 representing sperm cells.

M0 was represented by 945 cells and 87 significant marker genes, of which the top 43 markers have no reciprocal best hit in *C. elegans*. The 39^th^-ranked marker gene, however, had a top BLAST hit to *C. elegans swm-1*, which serves as an inhibitor of sperm activation within the male gonads^66,67^. Most of the top marker genes were Kunitz/Bovine pancreatic trypsin inhibitor domain proteins, and 14 of the 87 marker genes contained this InterPro domain (*P* = 5.1×10^-13^; **Supplementary Table S2M**). In the plant parasitic nematode *Bursaphelenchus xylophilus*, two such Kunitz effector proteins were found to be most highly expressed in the testis^68^. Marker genes from M0 were also significantly enriched for secretion in male ESPs in *A. ceylanicum*^12^ (*P* < 10^-15^). In a previous study, the “out of testis” hypothesis, which postulates that the testis represents a special environment facilitating the formation of novel genes, was strongly supported for nematodes including *C. elegans* and *Pristionchus pacificus*^69^. The cluster M0 markers were enriched for *Ancylostoma*-specific genes (*P* = 3.5×10^-5^), and 9 of the 95 MEROPS class I02 Kunitz-type protease inhibitors were among the top 19 markers for M0 (*P* < 10^-15^), eight of which came from I02 subclasses that were *Ancylostoma*-specific, providing further support for the speciation of these novel marker genes in *Ancylostoma*. Based on all of this evidence the M0 cell cluster is annotated as testis.

Two additional male-specific clusters (clusters M8 and M3) were likely to represent the two remaining male reproductive tissues in *A. ceylanicum*^59^: the cement gland and seminal vesicle. For M8 (129 cells and 86 marker genes) only four of the marker genes have reciprocal best BLAST hits to *C. elegans*, representing a significant depletion compared to random selection (*P* = 8.5×10^-17^), suggesting that this male-specific reproductive tissue was not conserved in *C. elegans*. The marker genes were strongly enriched for detection in both male and female ESPs^12^ (*P* < 10^-15^), with female expression levels for these markers only being high in cluster F0, the egg/embryonic tissue (see above). A cysteine-rich secretory proteins/antigen 5/pathogenesis-related 1 (CAP) domain was identified in 58 of the 86 marker genes for this tissue (*P* < 10^-15^; **Supplementary Table S2N**), and these were represented among the top secreted markers. Combining all this information supports the notion that this tissue was the cement gland. Overall, the secretory marker genes may represent a cocktail of proteins secreted from the cement gland with secreted sperm, in order to protect the sperm from the host environment. Their expression in eggs and embryos may represent that they play the same role in protecting those tissues from the host environment.

M3 (531 cells and 40 marker genes) was a male-specific cell type. Since we have relatively confidently assigned clusters for the sperm, seminal vesicle and cement gland, by the process of elimination, the remaining male reproductive tissue that we expect to observe with a lot of cells was the seminal vesicle. The 40 marker genes (**Supplementary Table S2O**) were very male-specific both within this dataset, and across the life cycle^11^. The top markers include many *Ancylostoma*-specific genes with unknown function, for which we were unable to deduce putative function based on lack of sequence similarity across the many reference databases used. Furthermore, to our knowledge there are no published reports identifying seminal vesicle-specific marker genes in nematodes. Here, we identify several transferases, phosphatases and glutamate carrier proteins as being markers for this potential seminal vesicle tissue in *A. ceylanicum*.

**Fig 8** shows a schematic (**Fig 8A**) of all the described male tissues, supported by structural observations by confocal microscopy (**Fig 8B**). In males, the testis was observed as a long coiled tubular structure in the middle region of the worm and connected to the seminal vesicle. The posterior end of the seminal vesicle connected to the cement gland which connected to the cloaca. However, no specific marker for seminal vesicle were identified, but its structure aligns with description in previous literature^59^. Two-photon microscopy revealed the copulatory spicules as prominent, rigid structure extending posteriorly into the cloaca and connecting to the ejaculatory duct. These essential reproductive structures facilitate mating by probing for the vulva opening and holding it open during ejaculation to assist sperm transfer^70^ (**Fig 8C**). Structural detail of the cloaca and cement gland (green), as well as prominent copulatory spicules (orange) and an ejaculatory duct (white), all consistent with previously-described reproductive anatomy for adult male *A. ceylanicum*^59^. DAPI-stained confocal microscopy (**Fig 8D**) identified additional structural detail including potential muscular structure in the adult male tail.

**Fig 8:**
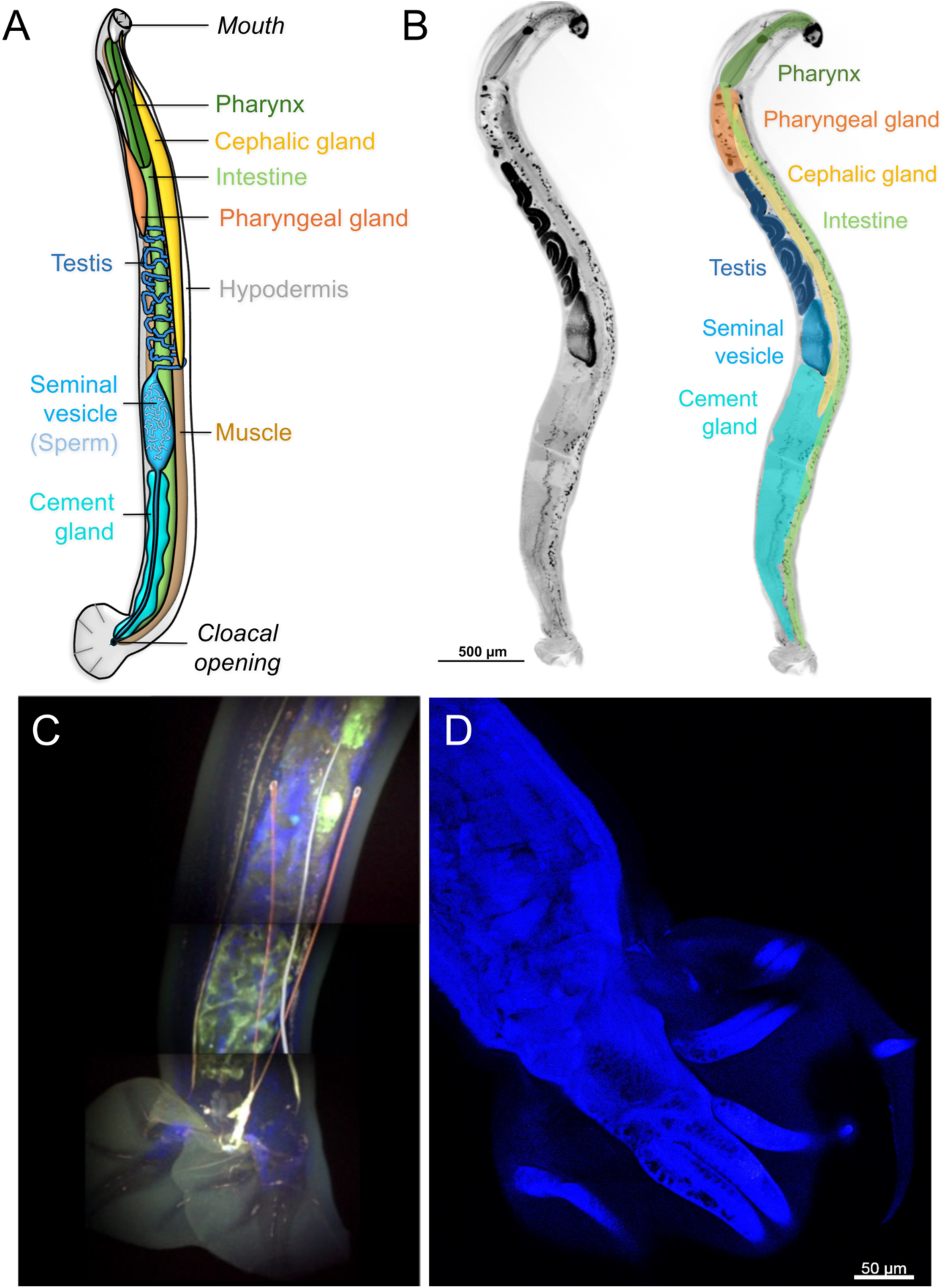
Microscopy of adult male *A. ceylanicum* worms indicating major tissues of interest. (**A**) Schematic representation of a male adult worms with major tissues indicated. (**B**) Confocal image of an adult male *A. ceylanicum* worm stained with BB, acquired at 20x magnification showing distinct anatomical features include the pharynx, pharyngeal gland, cephalic gland, intestine and reproductive organs including the testis, seminal vesicle and cement gland. (**C**) Two-photon microscopy image showing detail of the posterior region of an adult male *A. ceylanicum* worm, including the cloaca (rounded structures at the bottom), the cement gland (green), spicules (thin orange structures), and ejaculatory duct (thin white structures). Structures are inferred based on known anatomy^59^. (**D**) Confocal microscopy with DAPI staining of the posterior region of an adult male *A. ceylanicum* worm, showing additional structural details.

## Conclusions

In this study, we present the first single-cell transcriptomic atlas of a parasitic nematode, the adult male and female *Ancylostoma ceylanicum*, a human hookworm. The goal was to capture the heterogeneity of cells and annotate them at a tissue level. We confidently characterized 10 distinct cell types in females and 13 in males, including both shared non-reproductive tissues (pharynx, intestine, cephalic and pharyngeal glands, muscle, and hypodermis) and sex-specific reproductive tissues using multiple lines of evidence, including marker gene analysis, functional enrichment, comparative genomics with *C. elegans* and *A. suum.* As a validation for the cell type annotation, we performed FISH for a marker intestinal gene, and a high-resolution confocal microscopy for all the tissues/organ systems.

The analysis provided novel marker genes for specific tissues including the cephalic and pharyngeal glands, distinguished by the expression of CAP domain proteins, ASPs, TIMPs, and ShKT domain proteins, for which molecular studies have not previously been possible. We also identified distinct biology between the female and male pharynx, as well as distinct transcriptional characterization of tissues such as the seminal receptacle and the cement gland, which have been previously primarily only described anatomically in the literature for any nematode.

With this dataset, the resolution setting of 0.2 that was used appeared to be optimal in order to confidently distinguish and annotate relevant tissues. Throughout the analysis, there was just one instance in which clusters needed to be split to distinguish biologically relevant tissues (for the male muscle, hypodermis and neuronal clusters), and one instance in which clusters were merged (for the male pharynx) to more robustly define marker genes. We sought to provide the most comprehensive and useful biological output from this analysis, but any tissue of interest could be further subclustered and studied in more detail in future research. That was not the goal of this analysis, since performing the broad functional annotation of the major clusters is a substantial undertaking due to the lack of available such resources for parasitic nematodes. Future datasets sequencing more cells and deeper sequencing coverage may better support further subclustering.

This comprehensive atlas represents a significant advance in understanding hookworm biology at the cellular level and provides a valuable resource for the parasitology community. As hookworm infections continue to affect hundreds of millions of people globally, these insights into tissue-specific gene expression and function may accelerate the development of more effective treatments, vaccines, and diagnostic tools to combat this neglected tropical disease.

## Acknowledgments

We thank the GTAC team at the McDonnell Genome Institute at the Washington University in St. Louis for scRNAseq data generation. We also acknowledge Wandy L Beatty at the confocal imaging facility at Department of Molecular Microbiology and Mark J. Miller from the In Vivo Imaging Core (IVIC) in the Department of Medicine, Division of Infectious Diseases at the Washington university in St. Louis for providing training and access to the confocal microscopy. This work was supported by the National Institute of Health – NIAID grant AI159450 to M.M.

## Author contributions

Conceptualization: MM, BAR

Data curation: SB, BAR, JM

Formal Analysis: SB, BAR, MM

Funding acquisition: MM

Investigation: MM

Methodology: SB, BAR, MM

Project administration: MM

Resources: MM

Software: JM

Supervision: MM

Validation: SB, BAR, MM

Visualization: SB, BAR, JM

Writing – original draft: SB, BAR, MM

Writing – review & editing: SB, BAR, MM

## Data Availability

All raw data is available from the Sequence Read Archive (SRA), BioProject #PRJNA1147629, with accessions for samples provided in **Table 1**. The R scripts used for analysis are provided for download from GitHub (https://github.com/Sbmaurya-24/scRNA-seq-Adult-A.ceylanicum-). All data for all genes including functional annotations, cluster marker statistics and gene expression values per cluster are provided in **Supplementary Table S1**, and all marker gene lists and associated annotation data, expression data, and enrichment statistics are provided in **Supplementary Table S2**.

## Supplemental information titles and legends

**Supplementary Table S1**: Functional annotations, cluster marker statistics and gene expression values per cluster for every *A. ceylanicum* protein coding gene.

**Supplementary Table S2**: Marker gene significance values, annotations, expression data and functional enrichment data (provided as an external MS Excel file).

**Supplementary Fig S1:** Subclustering of the male hypodermis cluster to identify potential separate tissues. (**A**) Subclustering of the male hypodermis cluster (cluster 4) produces 3 distinct clusters, which were defined at a “FindClusters” resolution of 0.2. The three male hypodermis subclusters were annotated as neuronal, muscle and hypodermal tissues, and cluster separately on the male-only cell layout (**B**) and the female+ male integrated cell layout (**C**).

## STAR Methods

### Hamster infection with *A. ceylanicum* infective L3

Hamster infection with *A. ceylanicum* infective L3 was performed following the established protocol^12^. 3-4 week-old golden Syrian hamsters were gavaged using the appropriate technique and were accomplished according to the guideline of the Care and Use of Laboratory Animals of the National Institutes of Health (NIH). Freshly isolated infective *A. ceylanicum* L3 larvae were counted under the microscope and 80-100 L3 were suspended in 200ul of PBS per Eppendorf tube. A curved 18G needle was used to administer the infective L3. The hamsters manually restrained by gently grasping the loose skin at the scruff of the neck with the thumb and forefinger to immobilize the head and torso. Using the needle position adjacent to the hamster, the distance from the tip of the nose to the last rib on the left side was carefully measured. Subsequently, with the hamster’s nose pointed upwards, the gavage needle was introduced into the mouth over the tongue, directed towards the esophagus on the left side of throat, and gently pressed against the back of the mouth to facilitate swallowing. The needle was then slowly advanced to the premeasured distance, ensuring the accurate positioned, approximately 200ul of the larvae suspension was gently dispensed into the hamster’s stomach. Following the complete administration of the inoculum, the gavage needle was cautiously withdrawn, and the hamster was returned into the cage. Observation of gavaged hamsters for any sign of respiratory distress or bleeding the from the mouth or nose was conducted for a minimum of 5 minutes before returning them to the animal holding room.

### Isolation of adult *A. ceylanicum* from hamster intestine

To harvest the *A. ceylanicum* female and male adult worms, two hamsters were euthanize using appropriate protocol, equipment, and agents at twenty and twenty five days of post-infection (dpi) respectively with 80-100 infective iL3. Compressed CO2 gas in cylinders was used as a source of carbon dioxide, allowing for the controlled inflow of gas to the induction chamber. Hamsters were placed in the chamber and introduced 100% CO_2_ at a fill rate of 70% displacement of the chamber volume per minute with Co_2_, added to the existing air in the chamber, enabled the rapid and induction of unconsciousness with minimal distress to the hamsters. Each hamster was monitored for sign of unconsciousness, including lack of respiration and faded eye color. Confirmation of death was achieved by verifying the cardiac and respiratory arrest along with fixed and dilated pupils. Following euthanasia, the small intestine was removed, longitudinally cut and open. Adult worms were recovered from the intestine, collected, and washed thoroughly with 1x PBS containing 5x antibiotics/antimycotics and collected in the RPMI media with 10 % FBS containing the 5x antibiotics/antimycotics.

### *A. ceylanicum* adult worm single cell dissociation

Single cell dissociation was performed following the protocol^14^, Briefly, 10 female and 10 male *A. ceylanicum* adult worms were aliquoted in 1.5 ml microcentrifuge tube in triplicate and quadruplicate respectively. The adult worms were washed thrice with1ml of L15 medium (Gibco™ Leibovitz’s L-15 Medium, no phenol red). Worm’s pellets were resuspended in 200 μL of SDS-DTT solution (200 mM DTT, 0.25% SDS, 20 mM HEPES, pH 8.0, 3% sucrose) diluted in 1:4 in Leibovitz’s L-15 medium without phenol red and placed on a rocker for 8 minutes. The adult worms were washed with L-15 medium multiple times (a total of five times) to remove the residual SDS-DTT and until no smell was present. After washing worm’s pellet was resuspended in 200μl solution of pronase from S. griseus at a concentration of 15 mg/mL in L-15 medium. The reaction was monitored and pipetting continuously for approx. 30 minutes was done. The reaction was stopped when most of the adult worms were broken open and single cells were visible. The dispersed cells, remaining worms and debris were pelleted by centrifugation on 1000 rcf for 6 min at 4°C. The pellet was resuspended in a solution of cold L-15 medium containing 10% FBS and centrifuge 5 s at 1000 rcf to separate remaining large debris and single-cells. The top approx. 900 µl cells suspension was filtered using 70 μm cell strainers to separate cells from large debris from each tube. The filtrates were pooled, and cell count estimation was done using a standard hemocytometer.

### 10x Genomics preparation and sequencing

Isolated cells were prepared for GEM generation and barcoding. cDNA was prepared after the GEM generation and barcoding, followed by the GEM-RT reaction and bead cleanup steps. Purified cDNA was amplified for 11-13 cycles before being cleaned up using SPRIselect beads. Samples were then run on a Bioanalyzer to determine the cDNA concentration. GEX libraries were prepared as recommended by the 10x Genomics Chromium Single Cell 3’ Reagent Kits User Guide (v3.1 Chemistry Dual Index) with appropriate modifications to the PCR cycles based on the calculated cDNA concentration. For sample preparation on the 10x Genomics platform, the Chromium Next GEM Single Cell 3’ Kit v3.1, 16 reactions (PN-1000268), Chromium Next GEM Chip G Single Cell Kit, 48 reactions (PN-1000120), and Dual Index Kit TT Set A, 96 reactions (PN-1000215) were used. The concentration of each library was accurately determined through qPCR utilizing the KAPA library Quantification Kit according to the manufacturer’s protocol (KAPA Biosystems/Roche) to produce cluster counts appropriate for the Illumina NovaSeq6000 instrument. Normalized libraries were sequenced on a NovaSeq6000 S4 Flow Cell using the XP workflow and a 50x10x16x150 sequencing according to manufacturer protocol. A median sequencing depth of 50,000 reads/cell was targeted for each Gene Expression Library. The 10X Genomics Cell Ranger (version 7.1.0) pipeline was used to perform demultiplexing, barcode processing, and reads mapping to the *A. ceylanicum* genome^9^, using genes defined in using our improved genome annotation^12^.

### scRNA-seq bioinformatics analysis

Acey adult female and male data was downloaded from the cell ranger pipeline and read the data using Read10X () function. The resulting cell matrix was converted to a Seurat^18^ (v 4.4.0) object. Seurat commands and scripts were used for analysis below, unless otherwise indicated. The data was filtered to include only genes that were detected in a minimum of 3 cells and to encompass cells with at least 100 detected genes (i.e. min cells = 3, min features = 100). The SCTransform was used to normalization and variance stabilization and mitigate the technical noise and leading to the more accurate and interpretable downstream analyses. Before removing doublets, principal component analysis (PCA) was performed using 20 dimensions and was used as input for dimension reduction via UMAP, and clustering (‘FindNeighbors’ and ‘FindClusters’) at resolution 0.1. The DoubletFinder (v2.0.4)^71^ package was used to detect the potential doublets in the Seurat object with the estimated pN=0.25, pK=0.03 and expected doublet rate 7.5%.

After filtering out the doublet cells, data was normalized again using SCTransform for downstream analysis. Principal component analysis (PCA) was performed using 20 dimensions and was used as input for dimension reduction via UMAP, and clustering (‘FindNeighbors’ and ‘FindClusters’) at resolution 0.1 for both the female and the male samples (independently). There is no obvious or default resolution setting to define the number of clusters for the analysis, but our goal was to choose a cluster number that well-represented broad tissues that could be confidently annotated, without either grouping together transcriptionally distinct tissues, or over-dividing tissues into smaller groups, which would become difficult to distinguish and annotate. When performing clustering in Seurat^72^, clusters were numbered in order of the number of cells, by convention starting at cluster 0. In the interest of maintaining consistency between the raw data in the R object, and in repeatability of the analysis by future researchers, we have maintained this numbering throughout the manuscript. ‘FindAllMarkers’ was used to output MAST^73^ cluster marker statistics for each cluster, with significant markers parsed at a minimum Log_2_ fold change 0.25 (in cluster vs in other clusters), ‘Min. pct.1’ 0.25 (requiring at least 25% of cells in the cluster to have gene expression for the marker) and a *P* value ≤ 0.01. Male and female integrated analysis was performed using ‘IntegrateData’, with UMAP ran again using 20 principal components, for the purposes of visualization of cluster overlaps between the two samples and not for quantitative statistics. Subclustering of the male hypodermis cluster (cluster 4) was performed by separating the subset of cells assigned to that cluster (using the ‘subset’ function), using the same approach but a resolution of 0.2. Merging of the male pharynx clusters (clusters 6, 10 and 11) was performed by ‘WhichCells’ to define the cells belonging to the merged cluster, and ‘FindMarkers’ was used to compare those cells to all other male cells.

### Genomic annotation database construction

As described for the mapping procedure, a recently published improved annotation^12^ of the *A. ceylanicum* genome^9^. A functional annotation database was collected our the previous study^12^, including gene ontology^29^ classifications and InterPro functional domains^28^ from InterProScan v5.42^74^, and KEGG^27^ annotations from GhostKOALA v2.2^75^. Potentially secreted proteins were identified using SignalP v5.0^30^ for signal peptides and transmembrane domains (proteins with 2 or more transmembrane domains were not classified as secreted). PANNZER^25^ and Sma3s^26^ were used for protein naming. Normalized gene expression levels from RNA-seq samples collected across the *A. ceylanicum* life cycle^10,11^ and from the adult male *A. ceylanicum* intestine^19^ were also retrieved from remapping results described in our previous study^12^, along with adult male and adult female *A. ceylanicum* ESP proteomics results^12^. Heatmaps of gene expression data were generated based on the Z-scores of expression across the life cycle for each gene. All of these results for all *A. ceylanicum* genes are provided in **Supplementary Table S1.**

An analysis of protein sequence similarity to other species of interest was also collected from our previous study^12^, and included BLAST^21^ (NCBI blastp v2.7.1+) searches against protein sets from *Caenorhabditis elegans* (version PRJNA13758.WBPS16^22^) and *Ascaris suum*^23^ (PRJNA80881.WBPS17^22^). For these BLAST searches, only reciprocal best BLAST hits were considered (i.e., the proteins were each other’s top hit in each species), unless otherwise stated. BLAST results were used to identify 1:1 orthologs of tissue and cell-type marker gene lists from a previously-published *C. elegans* scRNAseq study^15^, and 1:1 orthologs of tissue-specific overexpressed genes from bulk RNA-seq from *A. suum*^13^. Protein conservation data across hookworms and human was also quantified using OrthoFinder^31^, as described in our previous study^12^. Briefly, protein sets were downloaded from WormBase Parasite^22^ (WBPS16) for *A. duodenale* (PRJNA72581) and *N. brasiliensis* (PRJEB511), and from Nematode.net^76,77^ for *N. americanus*^78^. The *A. caninum* protein annotation was downloaded from the Helminth Genomes Consortium paper^79^, and *Homo sapiens* (Human; GRCh38.103) was downloaded from Ensembl^80^. For each *A. ceylanicum* protein, the BLAST E value, alignment length, % identity and indication of reciprocal best hits is identified in **Supplementary Table S1**, along with OrthoFinder orthologous protein families, species membership and classification.

Gene Ontology (GO) term significant functional enrichment was performed with GOSTATS v2.70.0^81^ and for InterPro domains and KEGG pathways using WebGestaltR 0.4.6^82^ (FDR-adjusted P ≤ 0.05, minimum 3 marker genes). Functional enrichment was also tested among (i) orthologs of *A. suum* tissue-overexpressed genes, (ii) orthologs of tissue-enriched and cell type-enriched marker genes from *C. elegans*, (iii) proteins detected in adult female and male *A. ceylanicum* ESP proteomics, and (iv) *Ancylostoma*-specific, nematode-specific and human-conserved protein sets, according to the OrthoFinder^31^ testing described above. For these tests, negative binomial tests were used (MS Excel) for each set of marker genes of interest, requiring a minimum of 3 marker genes to be considered significant. For testing the orthologs of *C. elegans* and *A. suum* genes of interest, a background of only the *A. ceylanicum* genes with reciprocal best hits with these species were considered. FDR correction of resulting *P* values was performed within each data type. Significant enrichment results after FDR correction for all of these functional categories of interest are presented on the Supplementary Tables for each of the marker gene sets of interest.

For each gene, complete MAST marker gene statistics for each cluster are included in **Supplementary Table S1**, including average Log_2_ fold change values, the percentage of cells with detected expression of the gene (pct.1: across cells within the cluster, and pct2: across cells within all other clusters), and the unadjusted and FDR-adjusted *P* values for significance. Normalized expression values per gene per cluster calculated by SCTransform() are also provided in **Supplementary Table S1**, along with a heatmap visualization based on the Z-score of these expression values per gene. All of these MAST statistics are also provided for each marker gene in the additional Supplementary Tables.

### Confocal imaging

For confocal imaging, *A. ceylanicum* adult worms were harvested from hamster intestine at 29 dpi and washed with PBS containing 2x antibiotic/antimycotic five times. Both adult female and male worms were stained with Bisbenzimide (BB) at a concentration of 1mg/ml using 2µL in100µL of PBS for 3-4 hours. We subsequently stained some adult worms with phalloidin (Abcam, ab176753) 50 μg/mL in PBS for 1 hour at room temperature. Post staining, worms were washed thrice with PBS and fixed with 4% formaldehyde for 15 min at room temperature. Following fixation, the worms were again washed three time with PBS. The BB-stained adult male and female worms were mounted on the glass slide using ProLong Gold Antifade reagent (Invitrogen). Images of male and female worms were acquired with a Zeiss LSM880 laser scanning confocal microscope (Carl Zeiss Inc. Thornwood, NY) using excitation of a 405nm diode laser and photomultiplier tube detection wavelength of 415-470nm. ZEN black software (version 2.1 SP3) was used for image acquisition and for obtaining Z stacks and tiled images of entire worms. Images were acquired using a Plan-Apochromat 20X and 40X numerical aperture (NA) 0.8 objective.

#### Fluorescence in situ hybridization (FISH)

Worms were collected and washed with 1× PBS containing 0.3% TritonX-100. Fixation was performed in freshly prepared 4% formaldehyde in PBSTx (without calcium and magnesium) for 3 hours at room temperature with gentle agitation. Fixed worms were washed twice in PBSTx for 5 minutes each, dehydrated sequentially in 50% methanol for 5 minutes, followed by 100% methanol twice for 5 minutes each, and stored at −20 °C overnight. For rehydration, worms were incubated in 50% methanol for 5 minutes and then washed in PBSTx for 5 minutes. Worms were permeabilized with Proteinase K (10 µg/mL) in PBSTx containing 0.1% SDS for 10 minutes at 37°C. Samples were then fixed in 4% formaldehyde in PBSTx for 10 minutes at room temperature and washed twice in PBSTx for 5 minutes each. From this point, in situ hybridization was performed according to the protocol described by Molecular instrument (Los Angeles, California, USA)^83^ for the wholemount nematode larvae. Probes, buffers and hairpins were purchased from Molecular Instruments. Briefly Pre-hybridization was performed in prewarmed probe hybridization buffer at 37°C for 1 hour. Probe hybridization was carried out with a final probe concentration of 16 nM at 37°C for 16-24 hrs. Excess probe was removed by washing the worms using prewarmed probe wash buffer at 37°C (2× for 5 minutes and 2× for 30 minutes). This was followed by three washes in 5× SSCT at room temperature for 5 minutes each. For amplification, worms were pre-incubated in amplification buffer at room temperature for 30 minutes. Fluorescently labeled hairpins were snap-cooled using a thermal cycler (95°C for 90 seconds, cooled to room temperature in the dark for 30 minutes). Hairpin solutions were prepared in amplification buffer, and worms were incubated in this solution in the dark at room temperature for 20 hours. Excess hairpins were removed with sequential washes in 5× SSCT: 2× for 5 minutes, 2× for 30 minutes, and 1× for 5 minutes. Worms were then mounted on glass slides using ProLong Gold antifade mounting medium with DAPI (Invitrogen) and slides were stored at 4°C until imaging on Zeiss LSM880 confocal laser microscope. using excitation of a 405nm diode laser and photomultiplier tube detection wavelength of 415-470nm. ZEN black software (version 2.1 SP3) was used for image acquisition and for obtaining Z stacks and tiled images of entire worms. Most images were acquired using a Plan-Apochromat 20X numerical aperture (NA) 0.8 objective. To achieve better resolution of the sensory organs, images were acquired with Plan-Apochromat 40X NA 1.4 oil objective. Following acquisition, images were processed and exported using Zen 2.3 lite software.

### Two-photon microscopy

Adult *A. ceylanicum* worms were harvested from hamster intestine and washed five times with PBS containing 2x antibiotic/antimycotic and used for two-photon imaging. Live imaging of *A ceylanicum* male adult worms was performed using a custom-built two-photon (2P) microscope equipped with SlideBook Software (3i), a 1.0 NA 20x water dipping objective (Olympus) and a video-rate scan head (Thorlabs). Worms were maintained under physiological conditions during imaging by placing them in PBS on glass slide. Laser power was minimized to prevent thermal and photodamage. Intrinsic fluorescence signals were excited with a Chameleon Vision II Ti:Sapphire laser (Coherent) tuned to 850nm. Fluorescence emission detected simultaneously using four PMTs using dichroic filters (495, 540, and 595nm) to separate second harmonic signal from the tegument and auto fluorescence emitted from internal structures. Images were acquired as 41 consecutive 2 μm z-steps with image dimensions of 1024x1024 pixels at 0.5 μm/pixel. Multi-dimensional data was rendered in 3D using Imaris 9.9 (Biplane) and tiled image volumes were used to analyze the morphology of the cloaca

