## Supplementary Information for "A single-cell transcriptome atlas of adult male and female human hookworm *Ancylostoma ceylanicum*"

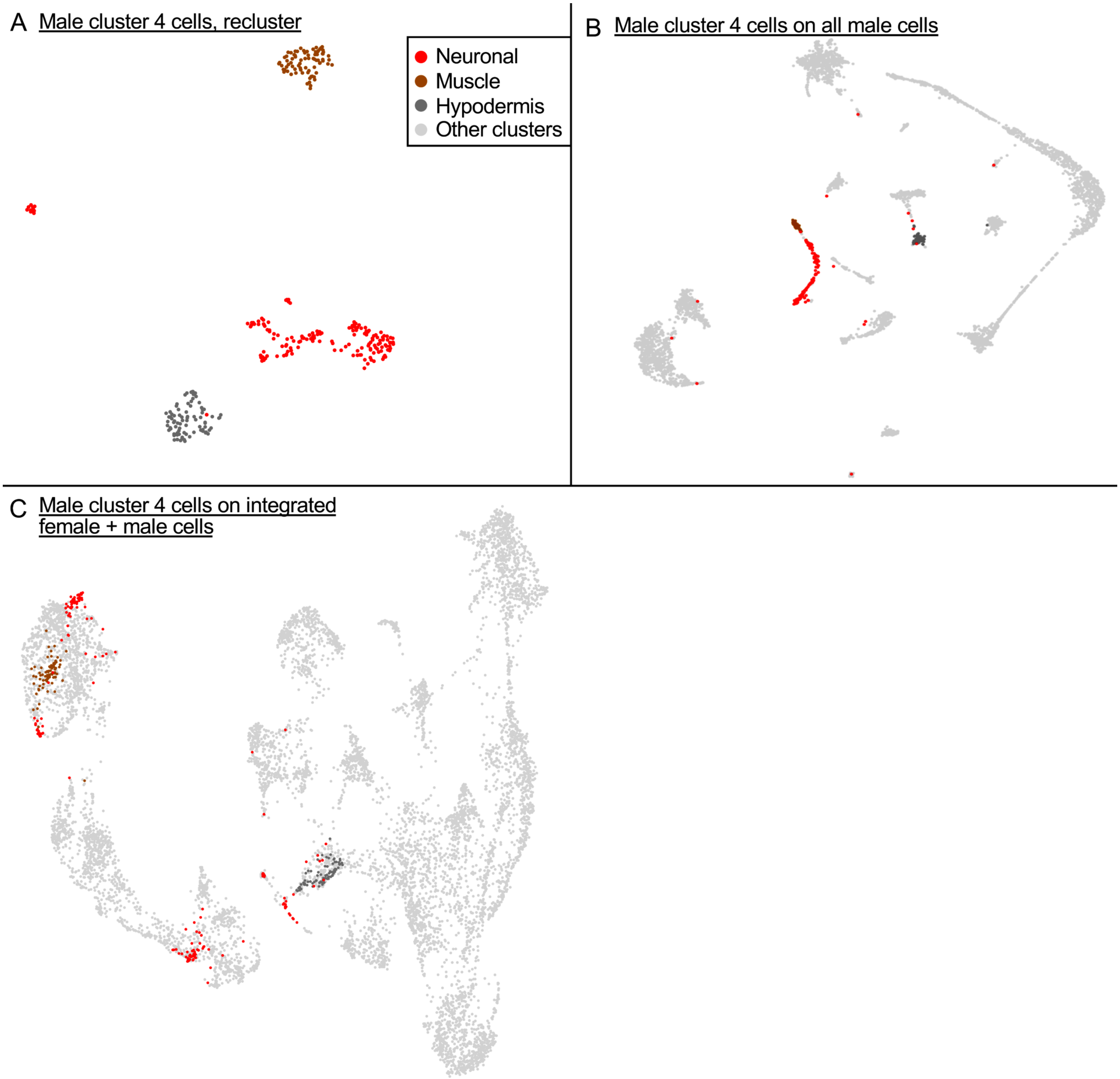


**Supplementary Fig S1:** Subclustering of the male hypodermis cluster to identify potential separate tissues. (**A**) Subclustering of the male hypodermis cluster (cluster 4) produces 3 distinct clusters, which were defined at a "FindClusters" resolution of 0.2. The three male hypodermis subclusters were annotated as neuronal, muscle and hypodermal tissues, and cluster separately on the male-only cell layout (**B**) and the female+ male integrated cell layout (**C**).
